# Barking up the right tree: Univariate and multivariate fMRI analyses of homonym comprehension

**DOI:** 10.1101/857268

**Authors:** Paul Hoffman, Andres Tamm

**Affiliations:** School of Philosophy, Psychology & Language Sciences, University of Edinburgh, UK

**Keywords:** ambiguity, context, comprehension, semantic control, hub-and-spoke

## Abstract

Homonyms are a critical test case for investigating how the brain resolves ambiguity in language and, more generally, how context influences semantic processing. Previous neuroimaging studies have associated processing of homonyms with greater engagement of regions involved in executive control of semantic processing. However, the precise role of these areas and the involvement of semantic representational regions in homonym comprehension remain elusive. We addressed this by combining univariate and multivariate fMRI analyses of homonym processing. We tested whether multi-voxel activation patterns could discriminate between presentations of the same homonym in different contexts (e.g., *bark* following *tree* vs. *bark* following *dog*). The ventral anterior temporal lobe, implicated in semantic representation but not previously in homonym comprehension, showed this meaning-specific coding, despite not showing increased mean activation for homonyms. Within inferior frontal gyrus (IFG), a key site for semantic control, there was a dissociation between pars orbitalis, which also showed meaning-specific coding, and pars triangularis, which discriminated more generally between semantically related and unrelated word pairs. IFG effects were goal-dependent, only occurring when the task required semantic decisions, in line with a top-down control function. Finally, posterior middle temporal cortex showed a hybrid pattern of responses, supporting the idea that it acts as an interface between semantic representations and the control system. The study provides new evidence for context-dependent coding in the semantic system and clarifies the role of control regions in processing ambiguity. It also highlights the importance of combining univariate and multivariate neuroimaging data to fully elucidate the role of a brain region in semantic cognition.

## Introduction

Language is laced with ambiguity. Most words have multiple semantic interpretations whose relevance depends on context. Often the various uses for a word appear to share a common semantic core; this is known as polysemy. This is not the case for homonyms like *bark*, however. *Bark* can refer either to the sound of a dog or to the covering of a tree, but these meanings have no semantic properties in common; they just happen to share the same phonological and orthographic form. Only around 7% of English words are homonyms (Rodd et al., 2002) but because their ambiguity is so distinct and well-defined, they represent a valuable test case for investigating how semantic processing is influenced by context more generally.

Psycholinguistic studies indicate that when we process homonyms in natural language, both meanings are briefly activated and compete for selection (Duffy et al., 1988; Vitello & Rodd, 2015). This competition is typically resolved within a few hundred milliseconds and the most contextually-appropriate meaning selected to guide ongoing comprehension (Seidenberg et al., 1982). Neuroimaging studies have implicated the left inferior frontal gyrus (IFG) in this meaning selection process. In listening tasks, left IFG shows stronger activation for sentences when they contain homonyms (Mason & Just, 2007; Rodd et al., 2005; Zempleni et al., 2007) and homonyms elicit greater IFG activation than unambiguous words in lexical decision and semantic judgement tasks (Bedny et al., 2008; Grindrod et al., 2014; Whitney et al., 2009).

These findings are well-explained by the controlled semantic cognition framework, which accounts for semantic processing in terms of interactions between semantic representation and control systems, supported by distinct neural networks (Hoffman et al., 2018; Jefferies & Lambon Ralph, 2006; Lambon Ralph et al., 2017). On this view, IFG is involved in top-down control over the activation and selection of semantic knowledge represented elsewhere in the cortex. Homonyms are assumed to place greater demands on this system because they require retrieval and selection of the contextually appropriate meaning (Noonan et al., 2010). Different functions have been ascribed to different subregions within IFG. The more anterior portion (pars orbitalis or BA47) is thought to support controlled retrieval of semantic knowledge, when the required knowledge is weakly associated with the stimulus and hence not activated automatically by spread of activation (Badre et al., 2005; Badre & Wagner, 2007). In contrast, posterior IFG (pars triangularis or BA45/44) is implicated in resolution of competition between active lexical-semantic representations (Nagel et al., 2008; Thompson-Schill et al., 1997). At present it is unclear whether the engagement of IFG during homonym processing reflects one or both of these processes, because both subregions show similar responses to ambiguous words (for discussion, see Vitello & Rodd, 2015).

Left posterior temporal regions also show increased fMRI activation in response to semantically ambiguous words, frequently centred on the posterior middle temporal gyrus (pMTG) (Humphreys & Lambon Ralph, 2017; Rodd et al., 2005; Rodd et al., 2015; Zempleni et al., 2007). The interpretation of these effects is more contested. One view is that pMTG is involved in control processes similar to those ascribed to IFG (Jefferies, 2013; Noonan et al., 2013; Whitney et al., 2011). Other researchers have proposed that pMTG is involved in representation of lexical-semantic information (or access to these representations), which would also be taxed during homonym comprehension (Bedny et al., 2008; Lau et al., 2008; Tyler et al., 2013). One fMRI study used multiple priming of homonyms (e.g., game-dance-ball) to attempt to differentiate between these accounts (Whitney et al., 2011). Both IFG and pMTG were sensitive to variation in the control demands of the task but not to the number of meanings that were retrieved on each trial, which appears inconsistent with a representational account. However, since ambiguity-related pMTG effects are weaker and less spatially consistent than those in IFG (Vitello & Rodd, 2015), there is less certainty over the role of this region.

The studies described thus far have used univariate activation contrasts to implicate IFG and pMTG in homonym comprehension. It is generally assumed that contrasts of this type index differences in the degree to which stimuli engage the processes or representations supported by the region (Taylor et al., 2013). Therefore, IFG and pMTG regions may show greater activation for homonyms because their multiple meanings necessitate greater engagement of the semantic control processes supported by these regions. What about regions implicated in representation of semantic knowledge? Current theories implicate anterior temporal and inferior parietal regions in semantic representation (Binder & Desai, 2011; Lambon Ralph et al., 2017). These areas do *not* typically show increased activation for homonyms, suggesting that the presence of ambiguity does not place greater metabolic demands on brain regions that encode semantic knowledge. But does this mean that these regions are not involved in meaning disambiguation? This seems unlikely. Multivariate fMRI studies have shown that activation patterns in anterior temporal and inferior parietal cortex vary according to the semantic properties of words and objects, supporting the idea that these regions code information about semantic content (Bruffaerts et al., 2013; Devereux et al., 2013; Fairhall & Caramazza, 2013; Peelen & Caramazza, 2012). It is therefore likely that elements of this network change their multivariate response to homonyms depending on the currently-relevant meaning, even in the absence of mean activation changes. Despite the recent burgeoning of multivariate fMRI studies, this hypothesis remains untested.

In the present study, we used multi-voxel pattern analysis (MVPA) to investigate how neural activation patterns vary in homonym comprehension, focusing on three representational brain areas in addition to the control-related regions discussed earlier. Two of these regions, the left angular gyrus (AG) and left lateral anterior temporal lobe (ATL), have been particularly implicated in semantic representation at a multi-word level. These regions show increased neural responses to coherent combinations of words, for example coherent adjective-noun phrases (e.g., *loud car*) or meaningful sentences (Bemis & Pylkkänen, 2012; Graves et al., 2010; Humphries et al., 2006; Price et al., 2015). Both regions have therefore been associated with combinatorial semantic processing, i.e., the extraction of a global meaning from a series of words (Bemis & Pylkkänen, 2012; Price et al., 2015; Vandenberghe et al., 2002). Other studies have found that both regions respond to non-verbal conceptual combination, suggesting a multimodal role in integrating and combining concepts (Baron & Osherson, 2011; Humphreys et al., 2019). The combinatorial semantic processing associated with both AG and lateral ATL would seem to be critical to homonym comprehension, where it is necessary to integrate the homonym with prior context in order to determine the appropriate semantic interpretation.

In contrast, the ventral portion of the ATL (inferior temporal and fusiform gyri) is strongly implicated in multimodal semantic processing at the single word/concept level (Lambon Ralph et al., 2017). Ventral ATL shows robust activation to individual words as well as multi-word combinations (Humphreys et al., 2015) and a series of multivariate neuroimaging studies have shown that activation patterns in this region discriminate object properties and word meanings (Clarke & Tyler, 2014; Coutanche & Thompson-Schill, 2015; Peelen & Caramazza, 2012). Theoretical accounts hold that the ventral ATL acts as a semantic hub that binds and integrates different linguistic and perceptual elements of experience to form coherent concepts (Lambon Ralph et al., 2017; Patterson et al., 2007; Rogers et al., 2004). Some accounts of ATL function posit that its representations must be context-independent, in order for conceptual knowledge to generalise appropriately across contexts (Binney et al., 2012; Lambon Ralph et al., 2017). Other models suggest that it is advantageous for the hub to be sensitive to context, to make use of semantic information present in the distributional statistics of language (Hoffman et al., 2018). Empirically, however, the degree to which lexical-semantic representations in the ventral ATL are independent of context remains an unanswered question. Homonyms provide a useful test case here because the same lexical item takes on radically different meanings in different contexts.

Only one previous fMRI study has investigated neural representation of homonyms in the ventral ATL. Musz and Thompson-Schill (2017) presented participants with homonyms, embedded in sentences that primed either their dominant or their subordinate meanings. They then compared the neural patterns elicited by the same homonym in the two different contexts. In the ventral ATL, they found that the similarity between dominant and subordinate patterns was predicted by the polarity of the homonym (i.e., the degree to which the dominant meaning occurs more often than subordinate one in natural language). When the dominant meaning was much more common than the subordinate one, the patterns in ventral ATL were more similar to one another. One interpretation of this result is that ventral ATL representations *do* vary as function of homonym meaning, but that highly dominant meanings are activated to some degree even when they are irrelevant. This may have caused the dominant and subordinate patterns to resemble one another for highly polarised homonyms.

In the present study, we investigated neural responses to balanced homonyms in which neither meaning was highly dominant over the other. We used univariate and multivariate fMRI to investigate patterns of neural engagement and information coding across the semantic network during homonym processing. By combining these distinct sources of information about homonym processing, we aimed to assess (1) the degree to which IFG sub-regions and pMTG support the control and selection of meanings and (2) the degree to which activation patterns in AG, lateral and ventral ATL vary according to homonym meaning. Participants were presented with sequential word pairs in which one meaning of a homonym was primed (e.g., *tree-bark* vs. *dog-bark*). This allowed us to assess whether activation patterns in each brain region could successfully discriminate between the alternative meanings of each homonym. On other trials, the prime was related to neither meaning (e.g., *letter-bark*), allowing us to test whether activity patterns discriminated the presence of absence of a semantic relationship. We also manipulated the task participants performed. In semantic runs, they decided whether the two words were semantically related; in phonological runs, they made syllable judgements in which the meaning of the words was irrelevant. This allowed us to assess the degree to which processes engaged during homonym comprehension occur automatically, in the absence of an explicit comprehension goal.

## Method

### Participants

24 native English speakers took part in the study (17 female; mean age = 22.3; sd = 4.2). All were classified as right-handed using the Edinburgh Handedness Inventory (Oldfield, 1971) and none reported dyslexia or any history of neurological illness. All provided informed consent. Data from one participant was excluded because they performed at chance when making semantic judgements about the meanings of homonyms (all other participants scored >85%). Neuroimaging and behavioural data for all participants are available on the Open Science Foundation repository: https://osf.io/ut3f9/.

### Stimuli

We investigated semantic processing of ten target words. Five of these were homonyms with two distinct meanings (*bark, calf, cell, pupil, seal*). The other five were unambiguous words matched to the homonyms for word length, frequency and concreteness (*coal, menu, monk, poet, wizard*). Each target word was paired with 12 different primes (see Figure 1B and 1C for examples). Eight of these were selected to have a strong semantic relationship with the target while the other four were unrelated in meaning. For the homonyms, half of the related primes were related to each of the target’s meanings, allowing us to investigate activity associated with opposing meanings of the same word.

**Figure 1:**
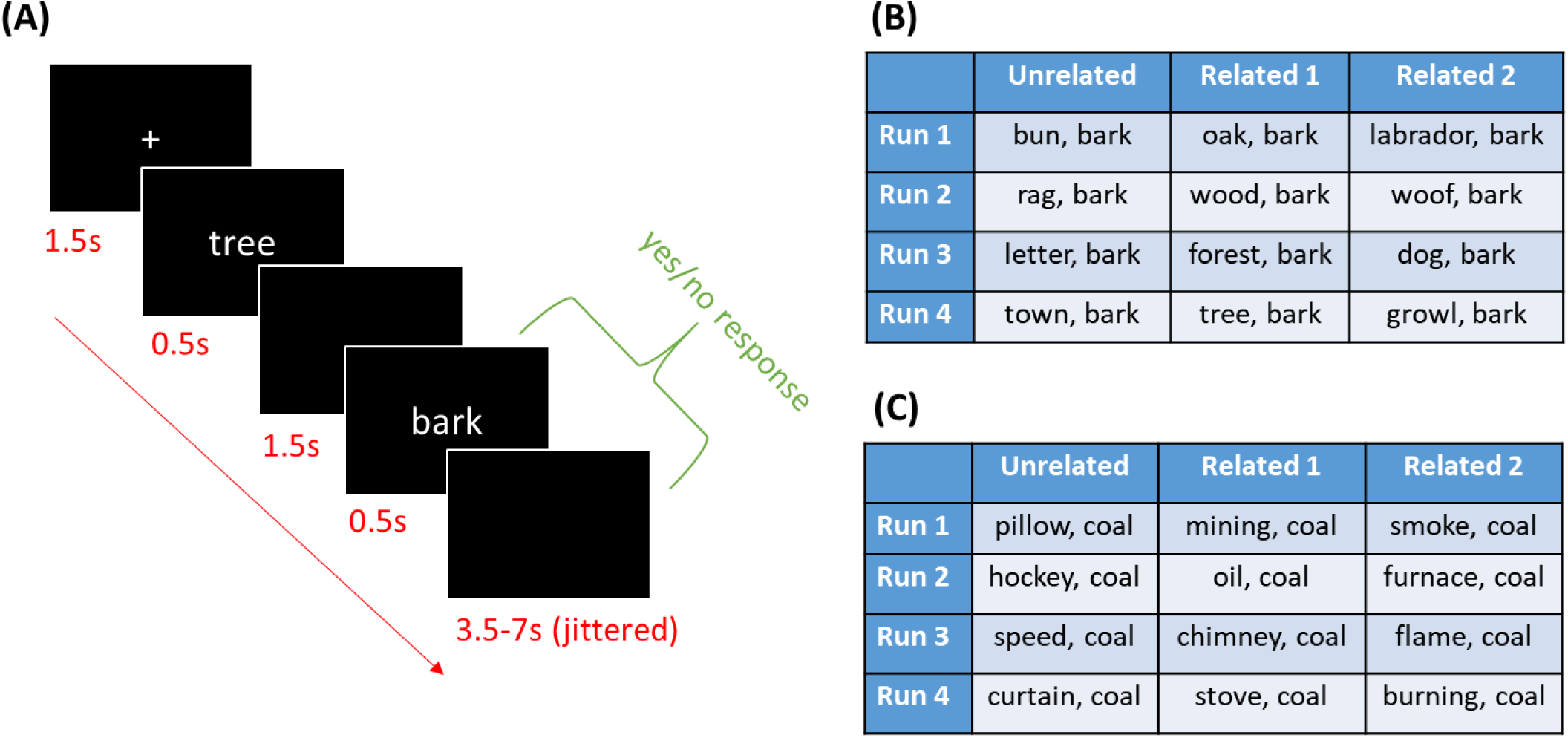
Experimental design. (A) Timeline for a single trial. (B) Prime-target pairs for a homonym target. (C) Prime-target pairs for an unambiguous target.

Semantic relatedness ratings were collected for all prime-target pairs from a group of 33 undergraduate students who did not take part in the main experiment. They rated the relatedness of the word pairs on a 5-point scale. Mean ratings by condition are reported in Table 1. The effects of relatedness over conditions was analysed with a 2 × 2 ANOVA (relatedness x ambiguity), which confirmed that related word pairs received higher ratings than unrelated word pairs and that relatedness did not differ between the Homonym and Unambiguous conditions. In addition, conditions did not vary in the frequency, concreteness and length of their primes (for full details, see Table 1). For the homonyms, care was also taken to ensure that the primes for each meaning had equally strong associations with the target. Stimuli were divided into four sets, for presentation in different scanning runs. In each set, each target appeared three times, once with an unrelated prime and twice with related primes. For homonyms, the two related primes in each set primed opposing meanings (see Figure 1B).

**Table 1:** Properties of primes in each condition

|  | Frequency | Concreteness | Length in letters | Length in syllables | Relatedness |
| --- | --- | --- | --- | --- | --- |
| Homonyms, Related | 4.14 (0.65) | 4.41 (0.62) | 5.55 (1.88) | 1.78 (0.83) | 4.49 (0.51) |
| Homonyms, Unrelated | 4.21 (0.62) | 4.30 (0.71) | 5.35 (1.93) | 1.55 (0.60) | 1.29 (0.20) |
| Unambiguous, Related | 4.15 (0.57) | 4.22 (0.84) | 5.98 (1.62) | 1.83 (0.75) | 4.58 (0.37) |
| Unambiguous, Unrelated | 4.01 (0.56) | 4.33 (0.56) | 6.05 (1.28) | 1.65 (0.49) | 1.22 (0.24) |
| Effect of Ambiguity | $F = 0.27$,<br>$p = 0.61$ | $F = 0.89$,<br>$p = 0.35$ | $F = 2.71$,<br>$p = 0.10$ | $F = 0.26$,<br>$p = 0.61$ | $F = 0.27$,<br>$p = 0.61$ |
| Effect of Relatedness | $F = 0.11$,<br>$p = 0.74$ | $F = 0.00$,<br>$p = 0.98$ | $F = 0.04$,<br>$p = 0.85$ | $F = 2.05$,<br>$p = 0.15$ | $F = 1938$,<br>$p < 0.001$ |
| Ambiguity x Relatedness interaction | $F = 0.75$,<br>$p = 0.39$ | $F = 0.69$,<br>$p = 0.41$ | $F = 0.17$,<br>$p = 0.68$ | $F = 0.03$,<br>$p = 0.86$ | $F = 1.19$,<br>$p = 0.28$ |

Table reports mean values per condition (standard deviations in parentheses) and results of 2x2 ANOVAs comparing conditions.
Frequency = Zipf frequency values from SUBTLEX corpus (Van Heuven et al., 2014);
Concreteness = values from (Brysbaert et al., 2014); Relatedness = Ratings of semantic relatedness of prime & target on a scale of 1-5.

### Procedure

Participants completed eight runs of scanning. In each run, they were presented with one set of 30 prime-target pairs. In half of the runs, they made semantic judgements about the prime-target pairs (are these words related in meaning?). In the other half, they made phonological judgements (do these words contain the same number of syllables?). The order of tasks was counterbalanced over participants. Manual responses were made using the left and right hands, with the mapping of these to response options counterbalanced over participants. Prior to entering the scanner, participants were warned that some of the targets had more than one meaning and that their primes could relate to either meaning. They were given brief definitions of the two meanings for each homonym (e.g., *Bark can mean the noise made by a dog or the covering of a tree*). We did this in order to ensure that participants were aware of both meanings and therefore likely to access them when primed appropriately during the main experiment. They practiced both tasks before entering the scanner.

The timeline for a single trial is shown in Figure 1A. Each trial began with a fixation cross presented for 1.5s. This was followed by the prime, which appeared for 0.5s. After a 1.5s delay, the target was presented for 0.5s. Participants were instructed to respond as quickly as possible upon seeing the target. Trials were separated by a mean inter-trial interval of 5s (jittered between 3.5s and 7s). Participants saw each set of trials once in the semantic task and once in the phonological task, with the order of presentation of the sets counterbalanced. Within runs, trials were presented in a different random order for each participant.

### Image acquisition and processing

Images were acquired on a 3T Siemens Prisma scanner using a 32-channel head coil. A dual-echo protocol was employed in which gradient-echo EPI images were simultaneously acquired at two TEs (13ms and 35ms) and a mean of the two echo series was computed during preprocessing (Halai et al., 2015). This approach improves signal quality in the ventral ATLs, which typically suffer from susceptibility artefacts (Ojemann et al., 1997). The TR was 1.8s and images consisted of 60 slices with a 100 x 100 matrix and voxel size of 2.4mm isotropic. Multiband acceleration with a factor of 2 was used and the flip angle was 74°. Eight runs of 158 volumes (284s) were acquired. A high-resolution T1-weighted structural image was also acquired for each participant using an MP-RAGE sequence with 0.8mm isotropic voxels, TR = 2.62s, TE = 4.5ms.

Images were preprocessed and analysed using SPM12. Preprocessing steps consisted of slice-timing correction, spatial realignment and unwarping using a fieldmap, normalisation of each participant’s images to a group template using Dartel (Ashburner, 2007) and finally transformation from the group template into MNI space. For univariate analyses, images were smoothed with a kernel of 8mm FWHM. Data were treated with a high-pass filter with a cut-off of 128s and the eight runs were analysed using a single general linear model. For each run, a single regressor modelled presentation of prime words, each with a duration of 0s, intended to capture brief neural processing associated with recognition and comprehension of the prime word. Targets were modelled with four regressors per run corresponding to the four experimental conditions (related-ambiguous, related-unambiguous, unrelated-ambiguous, unrelated-unambiguous). Each target was modelled with a duration of 2s. This longer duration was used to capture the neural processing associated with recognition of the target and with the subsequent semantic/phonological judgement. Covariates consisted of the six motion parameters and their first-order derivatives, as well as mean signal in white matter and CSF voxels.

Our main analyses focused on anatomical regions of interest (ROI) defined below. For univariate analyses, contrast estimates were extracted from these regions using Marsbar (Brett et al., 2002) and analysed with ANOVA. MVPA analyses are described below. We also conducted exploratory analyses at a whole-brain level, to ascertain whether experimental effects were present in other parts of the brain. Whole-brain univariate analyses were thresholded at a voxel level of *p*<0.005 and corrected for multiple comparisons at the cluster level using SPM’s random field theory (*p*<0.05 corrected for familywise error).

### Regions of interest

Our main analyses focused on left-hemisphere anatomical regions of interest (ROIs), selected a priori based on their involvement in semantic representation or control. These are shown in Figure 5A. Five of the six ROIs were defined using probability distribution maps from the Harvard-Oxford brain atlas (Makris et al., 2006), including all voxels with a >30% probability of falling within the following regions:

**Figure 2:**
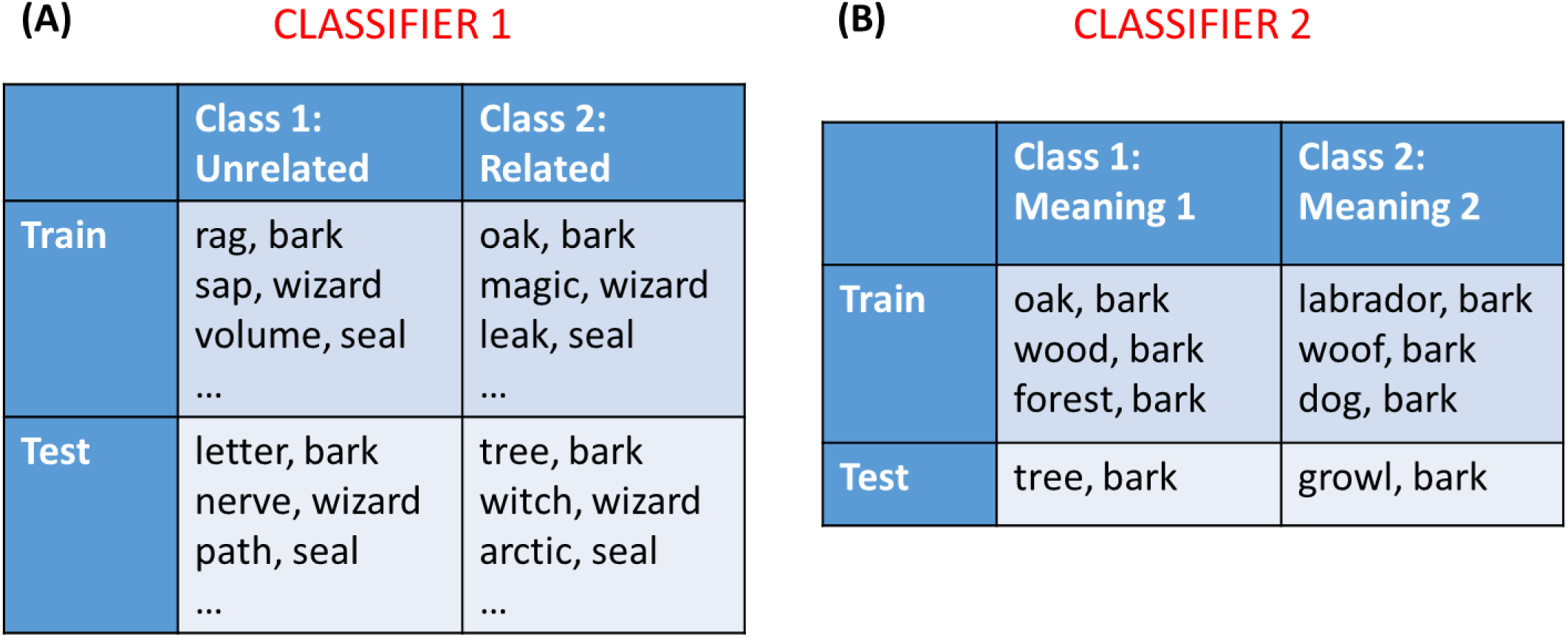
Illustration of multivariate classifier analyses. Figure shows example stimuli used to train and test each classifier in a single cross-validation fold. (A) Classifier 1 was trained to distinguish related from unrelated trials. All targets were included in train and test sets. Figure shows a subset of all trials that were included in the analysis. (B) Classifier 2 was trained to distinguish between the two meanings of each homonym. This analysis was performed separately for each homonym and the results averaged.

**Figure 3:**
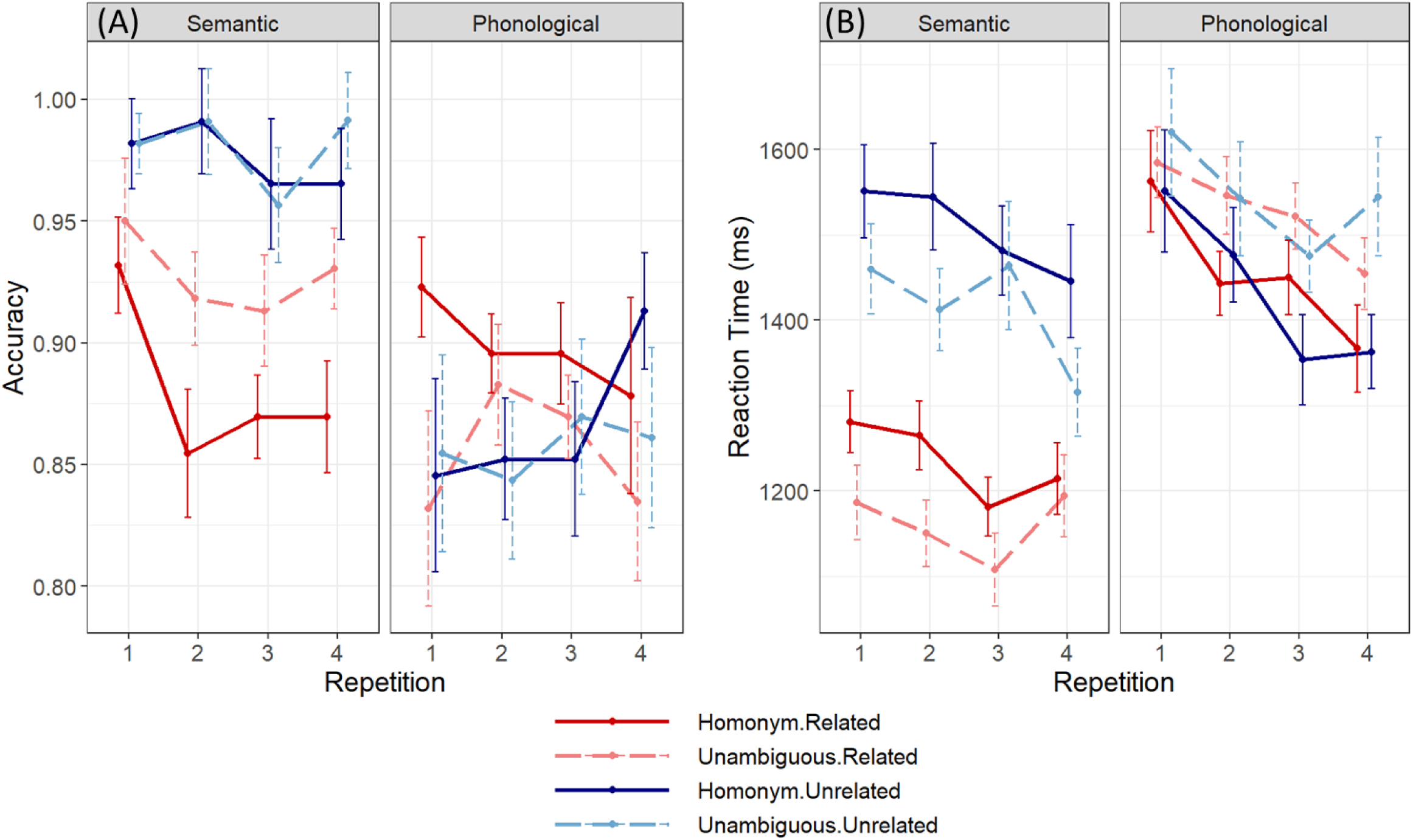
Behavioural performance. Bars indicate the standard error of the mean, adjusted to reflect the within-subject variance relevant for repeated-measures designs (Morey, 2008).

**Figure 4:**
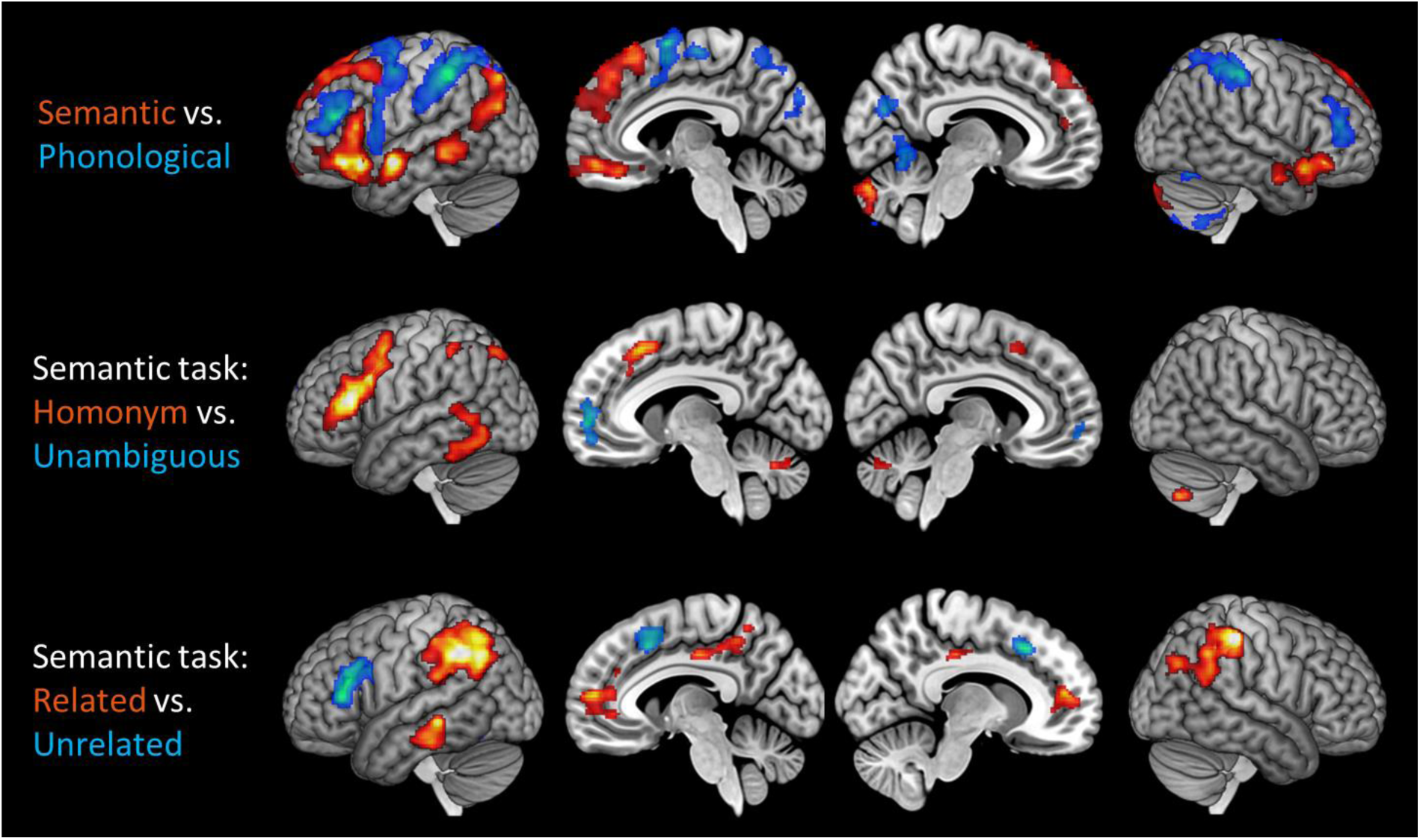
Whole-brain univariate activation contrasts. Images are shown at a voxelwise threshold of p<0.005, corrected for multiple comparisons at the cluster level.

**Figure 5:**
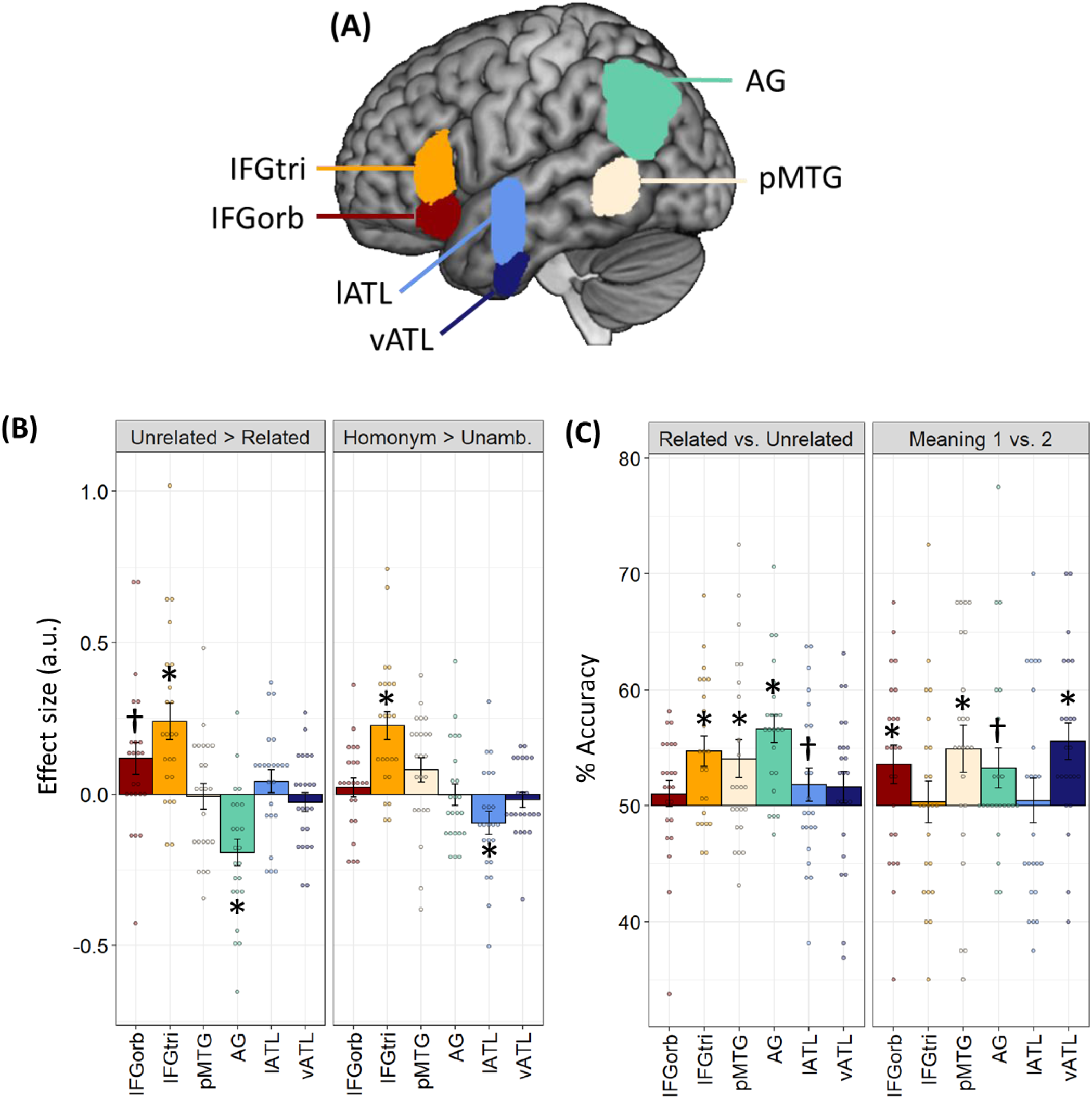
Region of interest results for the semantic task. (A) Locations of anatomically-defined ROIs. (B) Univariate contrasts of ambiguity and relatedness. (C) Classification accuracies for decoding analyses. Bars indicate between-subjects SEM; * indicates p < 0.05 following correction for multiple comparisons over all tests using the false-discovery rate (Benjamini & Hochberg, 1995). † indicates p < 0.05 with no correction for multiple comparisons.

IFGorb: the pars orbitalis region of inferior frontal gyrus, with voxels more medial than x=-30 removed to exclude medial orbitofrontal cortex (Hoffman, 2019)

IFGtri: the pars triangularis region of inferior frontal gyrus

pMTG: the temporo-occipital part of the middle temporal gyrus

Lateral ATL: the anterior division of the superior and middle temporal gyri

Ventral ATL: the anterior division of the inferior temporal and fusiform gyri

The final ROI covered the angular gyrus and included voxels with a >30% probability of falling within this region in the LPBA40 atlas (Shattuck et al., 2008). A different atlas was used in this case because the AG region defined in the Harvard-Oxford atlas is particularly small and does not include parts of the inferior parietal cortex typically implicated in semantic processing. The 30% inclusion threshold we used to define ROIs is also consistent with our previous work (Hoffman, 2019). Finally, it is worth noting that there are two common approaches to defining anatomical ROIs from probabilistic atlases, either the threshold-based method we used here or the maximum probability map method, in which voxels are assigned to the ROI if their probability exceeds that of falling into any other brain region (Eickhoff et al., 2006). To check that our ROIs were also consistent with the latter approach, we calculated the proportion of voxels in each ROI that were more likely to belong to its target region than to any other region. The vast majority of voxels in our ROIs met this criterion (mean over ROIs = 94%; range = 88-96%).

### Multivariate pattern analysis

For MVPA, normalised functional images were smoothed with a 4mm FWHM kernel (Hendriks et al., 2017). Each run was analysed with a separate GLM in which each of the 30 targets was modelled with a separate regressor (with a single regressor again modelling the presentation of primes). T-maps were generated for each target presentation and t-values from the voxels in each anatomical ROI were extracted for use in decoding analyses. The Decoding Toolbox (v3.997) was used for these analyses (Hebart et al., 2015).

Decoding analyses were performed separately on the four runs of the meaning task and the four runs of the phonological task. Classifiers were trained to discriminate between two classes of stimuli, using a support vector machine (from the LIBLINEAR library). To ensure independence of training and test data, we used a cross-validated leave-one-run out approach, in which the classifier was trained on data from three scanning runs and tested on the remaining run. The regularisation parameter *C*, which determines the classifier’s tolerance to misclassifications, was allowed to vary between 10^−4^ and 10^3^. The optimum *C* value for each training run was selected using leave-one-run-out nested cross-validation (Hebart et al., 2015).

Two forms of classification analysis were performed.

### Classifier 1: Related vs. unrelated trials

All targets were included in this analysis, coded according to their semantic relatedness, and the classifier was trained to discriminate related from unrelated trials (see Figure 2A). This analysis tested whether the neural patterns in each region reliably coded the presence or absence of a semantic relationship, irrespective of which particular word was being processed. The univariate analysis revealed that some regions showed differences in overall activation between related and unrelated trials. So that the classifier could not use these mean activation differences to discriminate the two classes, we mean-centred the activation patterns for each trial prior to classification (Coutanche, 2013).

To determine whether classification in each region was significantly better than chance, we used the permutation-based approach proposed by Stelzer et al. (2013), adapted for ROI data. For each participant, we trained and tested the classifier repeatedly on data in which the class labels had been randomly permuted. This process was repeated 100 times to provide an accuracy distribution for each participant under the null hypothesis (Stelzer et al., 2013). A Monte Carlo approach was then taken to determine the null accuracy distribution at the group level. Specifically, one accuracy value was selected at random from each participant’s null distribution and these were averaged to give a group mean. This process was repeated 100,000 times to generate a distribution of the expected group accuracy under the null hypothesis. Finally, the position of the observed group accuracy in this null distribution was used to determine a p-value (e.g., if the observed accuracy was greater than 99% of values in the null distribution, this would translate to a p-value of 0.01).

### Classifier 2: Meaning1 vs. meaning2 for homonyms

Only the related trials with homonym targets were used in this analysis. We took all related trials with the same homonym target and trained a classifier to discriminate which meaning was primed (see Figure 2B). We repeated this process for each of the five homonyms in turn and averaged the results to provide a mean accuracy for each participant. This analysis therefore tested for meaning encoding at a word-specific level. It determined whether the neural patterns in each region reliably coded which meaning of the homonym was accessed.

The Stelzer et al. method was again used to determine whether classification was significantly above chance at the group level. The difference here was that there were 35 permuted accuracy values for each participant (7 possible permutations for each of the 5 homonyms). When generating the group null distribution, each of the 100,000 iterations involved randomly selecting one permuted accuracy value for each homonym in each participant and averaging across these values. The position of the observed accuracy value in the null distribution was again used to determine statistical significance.

In addition to testing whether each ROI displayed above-chance classification, we also tested whether the performance of the classifier varied as a function of IFG sub-region (i.e., a classifier x region interaction). We used the Stelzer et al. permutation approach here as well. First, we calculated the observed value of the interaction by calculating the difference between accuracy of the two classifiers for IFGorb minus the same difference for IFGtri. To generate a group null distribution for this value, we generated 100,000 interaction values by repeatedly sampling from participant’s null distributions as described above. The position of the observed interaction value in this distribution was used to determine the p-value for the interaction.

Finally, to investigate classification performance outwith our ROIs, we conducted exploratory searchlight analyses over the whole brain. We used the same classification parameters as the ROI analyses, but applied these to a spherical searchlight of radius 12mm that was moved iteratively across the brain. This provided classification accuracy maps for each participant. To perform group-level inference on these maps, a permutation-based approach was again used. The SnPM13 toolbox was used to determine the null distribution (10,000 permutations, 8mm variance smoothing) and to threshold maps at a voxel level of *p*<0.005, correcting for multiple comparisons at the cluster level (*p*<0.05 corrected for familywise error). Individual accuracy maps were smoothed with a 4mm FWHM Gaussian kernel prior to group analysis.

## Results

### Behavioural performance

Effects of the experimental manipulations on response accuracy and RT were investigated. Because the same targets were presented multiple times (four times per target per task, albeit with a different prime on each occasion), the effect of target repetition was also investigated. Figure 3A shows mean accuracy as a function of condition and repetition. These data were analysed using a generalised linear mixed effects model with fixed effects of task, ambiguity, meaning relatedness and repetition, and their interactions. The model included random intercepts and slopes for participants and trial number was included as a covariate. Significance of fixed effects were assessed using a likelihood ratio test (Baayen et al., 2008); hence chi-squared statistics are reported.

The main effect of repetition did not reach statistical significance (*χ*^2^ = 7.77, *p* = 0.051), nor did repetition interact with any other factors. There was, however, a main effect of task (*χ*^2^ = 19.9, *p* < 0.001), as overall performance was better for the semantic task, and relatedness (*χ*^2^ = 21.4, *p* < 0.001), since more correct responses were given when prime and target were semantically unrelated. There were also significant interactions of task with both relatedness (*χ*^2^ = 26.7, *p* < 0.001) and ambiguity (*χ*^2^ = 5.8, *p* = 0.015). Separate analyses performed on each task indicated that relatedness influenced performance on the semantic task (*χ*^2^ = 15.7, *p* < 0.001) but not on the phonological task (*χ*^2^ = 0.4, *p* = 0.51). This was expected, since the relatedness of the word pairs was irrelevant for the phonological task. In contrast, ambiguity only influenced accuracy on the phonological task (*χ*^2^ = 4.3, *p* = 0.038).

Reaction time data are presented in Figure 3B. Analyses were performed on these data following log-transformation to reduce skew. Model structure was the same as for accuracy but in this case a linear model was fit, in line with the continuous nature of the reaction time data. The model indicated an effect of repetition (*F*(3,31.9) = 5.4, *p* = 0.004), as participants tended to become faster with greater familiarity with the tasks and stimuli. There were also main effects of task (*F*(1,21.5) = 16.6, *p* < 0.001) and relatedness (*F*(1,22.6) = 54.4, *p* < 0.001). On average, participants were faster to respond in the semantic task and when the prime and target were semantically related. The task manipulation interacted with relatedness (*F*(1,22.7) = 41.8, *p* < 0.001) and ambiguity (*F*(1,24.5) = 31.9, *p* < 0.001). Separate analyses for each task revealed that, for the semantic task, participants were faster to respond to related word pairs (*F*(1,22.2) = 54.9, *p* < 0.001), while no such effect was present in the phonological task (*F*(1,2233) = 0.005, *p* = 0.94). In the semantic task, participants were slower to respond to homonym targets (*F*(1,24.1) = 21.5, *p* < 0.001). Conversely, they were faster to respond to homonyms in the phonological task (*F*(1,25.5) = 15.2, *p* < 0.001). This result is consistent with previous studies showing that lexical ambiguity has a negative effect when people make semantic judgements but is beneficial in other lexical processing tasks (Hino et al., 2002; Hoffman & Woollams, 2015).

### Neuroimaging contrasts of task

Whole-brain contrasts for the semantic vs. phonological task are shown in Figure 4 (for peak activation co-ordinates, see Supplementary Table 1). The semantic task produced greater activation in a range of predominately left-lateralised cortical regions implicated in semantic processing, including IFG, anterior and posterior temporal regions and AG. Greater activation for the phonological task was observed in frontal and parietal regions associated with phonological processing, working memory and cognitive control. It therefore appeared that participants activated highly distinct neural networks for the two tasks. In addition, the behavioural data indicated that our experimental manipulations had markedly different effects in each task. For these reasons, from this point forward we analysed neuroimaging data from each task separately.

### Univariate effects in the semantic task

Whole-brain effects of relatedness and ambiguity in the semantic task are presented in Figure 4 (for peak activation co-ordinates, see Supplementary Table 2). As expected, greater activation for homonyms was predominately observed in left prefrontal cortex and posterior temporal cortex. Semantically unrelated trials also engaged left prefrontal regions to a greater extent, while more activation on related trials was found in a number of default mode network regions, including bilateral AG, posterior cingulate and ventromedial prefrontal cortex. We also tested whether the relatedness effect interacted with ambiguity. It did: the Related > Unrelated effect in default mode regions tended to be stronger for the homonym trials (see Supplementary Figure 1).

Effects of ambiguity and relatedness within our ROIs are shown in Figure 5B (Supplementary Figure 2 shows activations in each condition relative to rest). We first performed a 6 × 2 × 2 ANOVA (region x ambiguity x relatedness) on these data to determine whether the effects of our experimental manipulations varied across regions. They did: region interacted with both ambiguity and relatedness (*F*(5,110) > 13.2, *p* < 0.001). There was no three-way interaction between the factors (*F*(5,110) = 0.31, *p* = 0.90). We therefore tested for the effects of ambiguity and relatedness in each region separately, using t-tests. Significant effects, with and without correction for multiple comparisons, are highlighted in Figure 5B.

As shown in Figure 5B, greater activation on homonym trials was observed in IFGtri only. Although whole-brain analysis identified greater activation for homonyms in the area of pMTG, within our ROI the effect failed to reach significance (uncorrected *p* = 0.058). In contrast, lateral ATL showed less activation for homonyms than for unambiguous words. Divergent effects of semantic relatedness were also present. IFGtri showed greater activation for the more difficult unrelated trials; IFGorb showed a similar effect though it did not survive correction for multiple comparisons. AG, conversely, was more engaged during related trials. Finally, as we were particularly interested in discriminating between the roles of the two inferior prefrontal regions, we conducted a 2 × 2 × 2 ANOVA contrasting experimental effects in IFGorb vs. IFGtri. IFGtri showed larger effects of the ambiguity manipulation (ambiguity x region interaction: *F*(1,22) = 22.1, *p* < 0.001) and the relatedness manipulation (relatedness x region interaction: *F*(1,22) = 5.72, *p* = 0.03).

### Multivariate pattern analysis in the semantic task

Decoding accuracies for the MVPA classifiers are shown in Figure 5C. The Related vs. Unrelated classifier tested whether neural patterns reliably signalled the presence or absence of a semantic relationship, across all words. Three regions exhibited this coding at an above-chance level: IFGtri, pMTG and AG. No coding of trial status was found in IFGorb or in the lateral or ventral ATL. The Meaning 1 vs. 2 classifier tested whether neural patterns reliably distinguished between the different meanings of each individual homonym. Three regions displayed this word-specific neural coding: IFGorb, pMTG and vATL. This indicates that the neural patterns in these regions reliably signalled which meaning of the homonym was relevant to the trial, suggesting that these areas code the opposing meanings of homonyms differently. AG showed a weaker effect that did not survive correction for multiple comparisons.

These results suggest that, during the semantic task, different parts of the IFG coded different types of information about the stimuli. To determine whether this was the case, we used permutation testing to test for an interaction between classifier analysis and IFG sub-region. The interaction was significant (*p* = 0.011), confirming that the two IFG subregions code different information about each trial. IFGtri appeared to code semantic status (related or unrelated) at a general level, while IFGorb coded word-specific information about the relevant meaning.

The whole-brain searchlight for the Related vs. Unrelated classifier is shown in Figure 6. It is important to remember that, in the semantic task, participants made different motor responses on related vs. unrelated trials. Above-chance decoding was therefore observed in bilateral premotor and motor cortices, as well as in prefrontal, temporal and parietal regions associated with semantic processing. We also performed a searchlight analysis for the Meaning 1 vs. 2 classifier but no effects were found at a cluster-corrected threshold.

**Figure 6:**
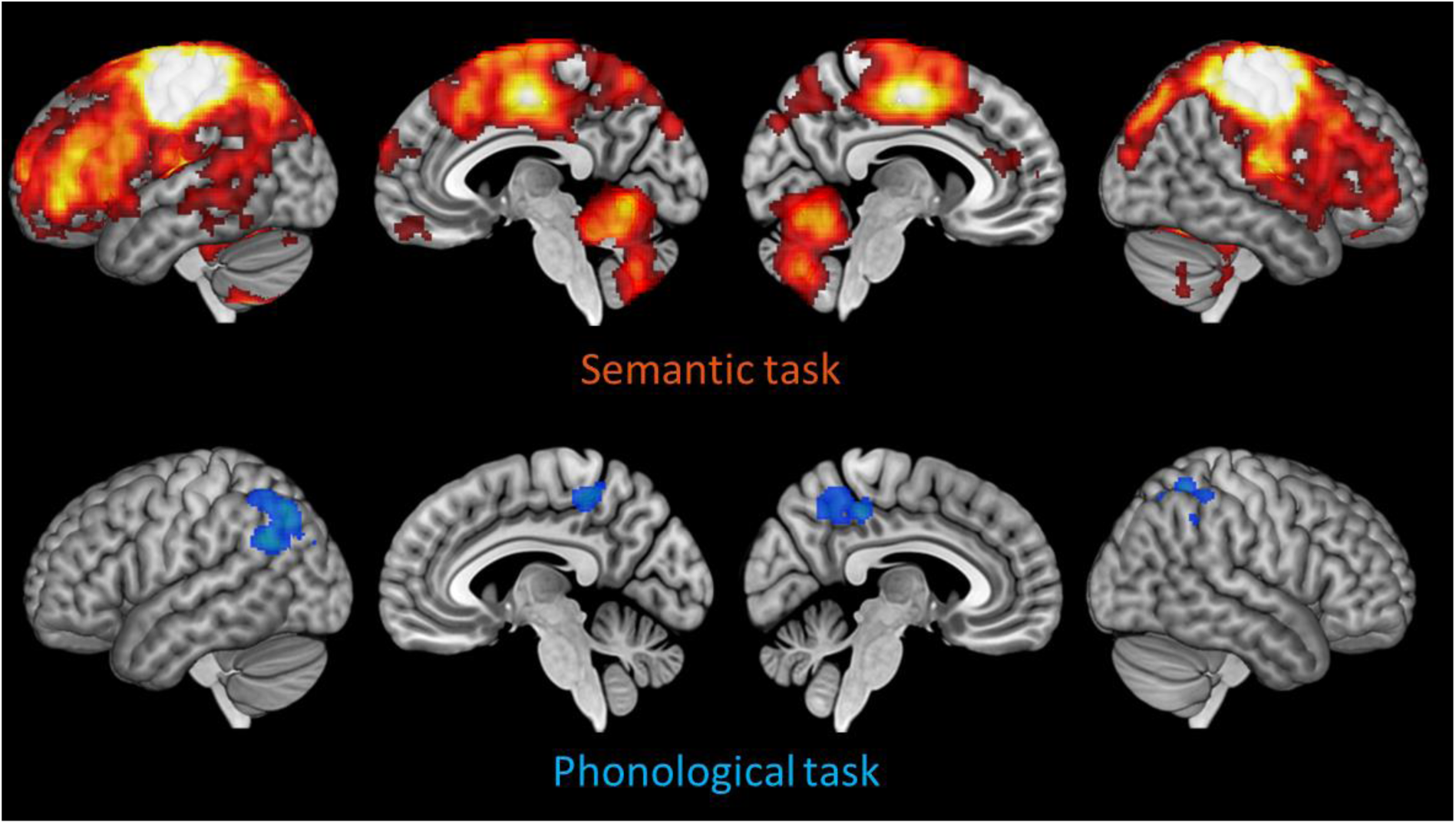
Whole-brain searchlight results for the Related vs. Unrelated classifier. Images are shown at a voxelwise threshold of p<0.005, corrected for multiple comparisons at the cluster level.

### Univariate effects in the phonological task

Whole-brain analysis for the phonological task revealed no effects of ambiguity. However, greater engagement for semantically related word pairs was found in bilateral AG, bilateral middle frontal gyrus and right IFG (see Supplementary Figure 3). Effects in ROIs are shown in Figure 7A. A 6 × 2 × 2 ANOVA (region x ambiguity x relatedness) identified an interaction between relatedness and region (*F*(5,110) = 2.76, *p* = 0.021) but no other interactions. When testing for effects in individual ROIs, significantly greater activation for related trials was found in AG. Greater activation for homonyms in IFGtri was observed but this effect did not survive correction for multiple comparisons.

**Figure 7:**
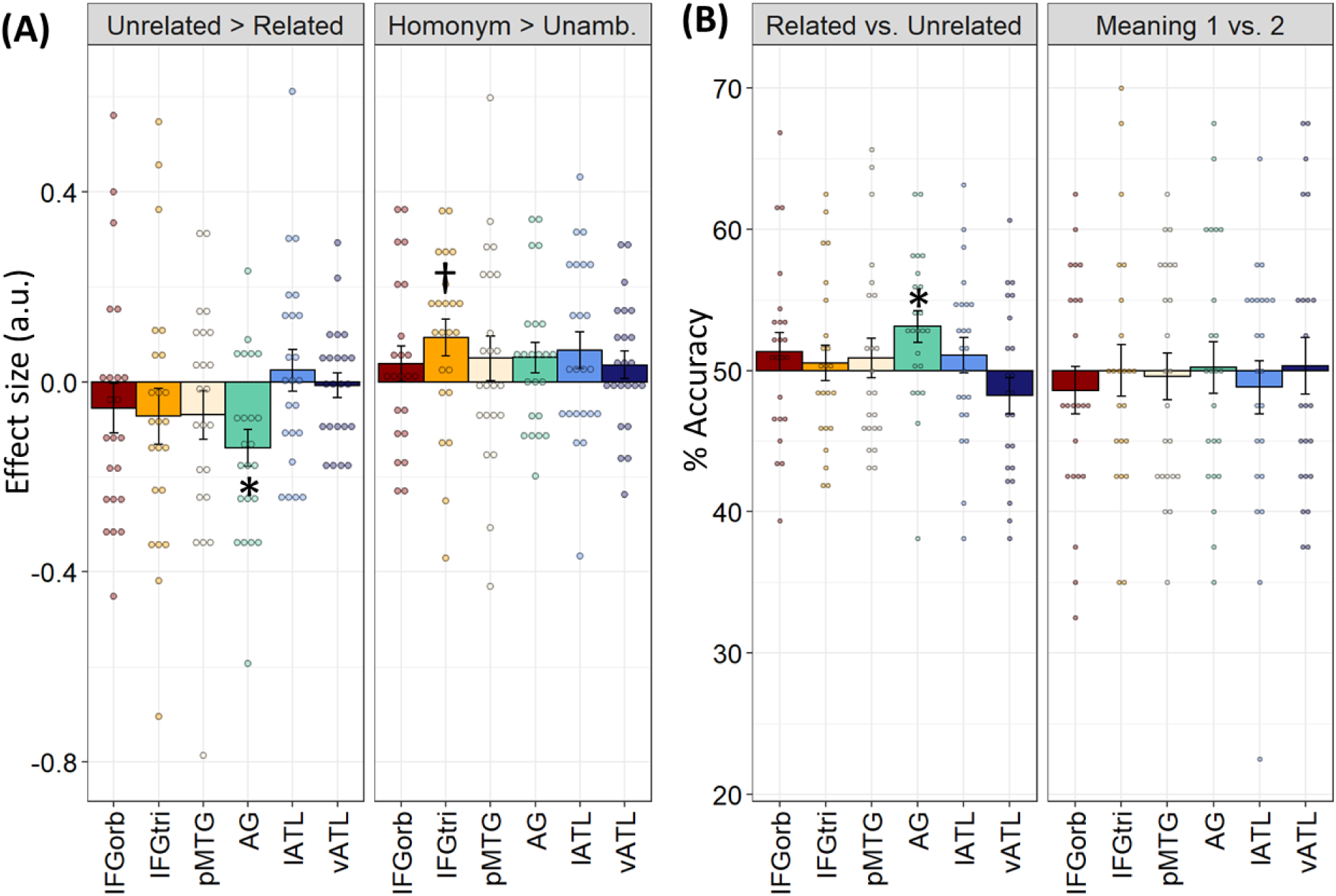
Region of interest results for the phonological task. (A) Univariate contrasts of ambiguity and relatedness. (B) Classification accuracies for decoding analyses. Bars indicate between-subjects SEM; * indicates p < 0.05 following correction for multiple comparisons over all tests using the false-discovery rate (Benjamini & Hochberg, 1995). † indicates p < 0.05 with no correction for multiple comparisons.

### Multivariate pattern analysis in the phonological task

Decoding accuracies for the MVPA classifiers are shown in Figure 7B. Patterns in AG showed above-chance decoding for the Related vs. Unrelated distinction, even though this distinction was irrelevant to the phonological judgements. Other ROIs showed no evidence of coding this information, even those that did decode this information during the semantic task. In the Meaning 1 vs. 2 classifier, no ROIs were able to discriminate between the different meanings of the homonyms.

The searchlight analysis for the Related vs. Unrelated classifier revealed significant decoding in bilateral parietal cortices and the precuneus (Figure 6). No significant clusters were identified by the Meaning 1 vs. 2 searchlight.

## Discussion

We used univariate and multivariate fMRI to investigate how different elements of the semantic neural network process the variable meanings of homonyms. Our principal finding was that various areas in left frontal and temporal cortices exhibited activation patterns that reliably predicted which of the homonym’s meanings was relevant to the trial. Previous studies have found that neural coding patterns vary according to the word being comprehended (as reviewed by Bruffaerts et al., 2019). Here, however, we found regions that displayed systematic variation in the patterns elicited by the *same* word in different contexts. This effect could occur because the region represents the word differently depending on which of its meanings is currently relevant. Alternatively, it could be that the two meanings place different processing demands on the region (e.g., one meaning is consistently more difficult to process than the other).

A distinct set of regions coded at a more general level for the semantic status of each trial. Importantly, multivariate stimulus coding was observed in the absence of univariate activation differences between trial types and vice versa. Our results are supportive of a broad distinction between regions that represent semantic knowledge and those that regulate and control its use. They also provide new insights into distinct roles played by different elements of the semantic network when processing ambiguity. In what follows, we first discuss interpretation of effects in individual regions before considering implications for more general accounts of the semantic system.

### Inferior frontal gyrus

A major aim of the study was to clarify the role of left IFG in resolving semantic ambiguity. Although the entire IFG region is implicated in semantic control, researchers have proposed a distinction between anterior IFG (IFGorb), which is thought to play a key role in controlling the activation and retrieval of knowledge from semantic memory (controlled retrieval), and the more posterior portion (IFGtri), which is involved in selecting task-relevant representations (Badre et al., 2005; Badre & Wagner, 2007; Nagel et al., 2008; Thompson-Schill et al., 1997). This basic distinction is reflected in computational models of semantic processing. In connectionist models of cognition, selection of task-relevant representations or responses is frequently achieved through representations of current goals that act to potentiate relevant processing units and inhibit irrelevant ones (Botvinick & Cohen, 2014). The most well-known instantiation of this approach is in models of the Stroop effect (Cohen et al., 1990), though models of semantic cognition have used similar methods (Dilkina et al., 2008; Plaut, 2002). However, Hoffman et al. (2018) argued that a different mechanism is required to mediate the controlled retrieval of conceptual information. Rather than using goal representations, their connectionist semantic network implemented controlled knowledge retrieval by allowing multiple retrieval cues to jointly constrain the settling of the network. This mechanism improved the network’s ability to detect associations between two concepts. Detailed discussion of these computational frameworks is beyond the scope of the present work. We mention them here, however, to highlight that both neuroimaging and computational studies support the general notion that semantic control functions involve separable processes for control of knowledge retrieval and competition resolution.

Despite this hypothesised functional dissociation between IFGorb and IFGtri in semantic processing, previous studies have provided little evidence for distinct roles in processing semantic ambiguity (Vitello & Rodd, 2015). In the present study, however, we found that these areas showed divergent profiles in both their levels of overall neural engagement and in the type of stimulus information coded in their neural patterns. In univariate analyses, IFGtri showed greater activation for homonyms compared with unambiguous targets but IFGorb did not. MVPA analyses also produced divergent results. In IFGtri, neural patterns reliably coded whether a semantic relationship was present. In contrast, IFGorb neural patterns coded item-specific semantic information, shifting their patterns of activity according to which meaning was currently relevant.

Results in IFGorb broadly support the assertion that this area supports controlled retrieval of semantic information, which is required when automatic stimulus-driven activity is insufficient to activate the necessary semantic representation (Badre & Wagner, 2007). The controlled retrieval account predicts greater activation on semantically unrelated trials, because these require a sustained retrieval effort in order to thoroughly search semantic memory and discount the possibility that a semantic relationship is present. IFGorb did exhibit this pattern, though only at an uncorrected statistical threshold. No effect was observed in the phonological task, since a controlled search for meaning is not engaged when participants are not motivated to process the stimulus at a semantic level.

Importantly, our MVPA results also indicate that IFGorb was engaged in meaning-specific retrieval processes, since the neural patterns in this region varied depending on the particular meaning that was relevant to the trial. This could have occurred because some meanings are consistently more difficult to retrieve than others and hence produce greater activation in this area. Or it could be that different meanings require the use of different cues or strategies to facilitate retrieval, resulting in distinct activation patterns. In contrast, patterns did not vary according to the required behavioural response (related vs. unrelated), suggesting that IFGorb is not involved in using semantic information to determine a behavioural response.

We did not observe greater univariate activation in IFGorb when participants processed homonyms. This is surprising as these words might have been expected to place greater demands on controlled retrieval in order to activate the correct aspect of meaning. There are two potential explanations for this. First, the appropriate meaning was primed prior to presentation of the homonym itself, which may have reduced the need for control. Second, we used balanced homonyms in which both meanings were similarly frequent in language and therefore both relatively accessible in the semantic system. Other studies that have used biased homonyms have found greater IFGorb activation when subordinate (infrequent) meanings are retrieved (Mason & Just, 2007; Whitney et al., 2009), in line with a greater need for controlled retrieval (Hoffman et al., 2018; Noonan et al., 2010).

In IFGtri, greater activation for unrelated trials and for homonyms is consistent with the more general selection and competition resolution mechanisms attributed to this region. On semantically unrelated trials, participants must reject and inhibit any irrelevant semantic information accessed as participants search for a meaningful connection between the words. Greater IFGtri engagement for homonyms is also readily explained in terms of competition between their two alternative meanings. In addition, related trials for homonyms induce additional competition between potential response options, since any activation of the currently-irrelevant meaning would direct participants towards making an “unrelated” response (Pexman et al., 2004).

MVPA results indicate that activation patterns in IFGtri were attuned to the correct behavioural response for the trial, but not to which word-specific elements of meaning were retrieved. This result suggests that IFGtri is less closely involved in processing the semantic properties of the stimuli per se, and more in using the semantic information to determine an appropriate behavioural response. Thus, overall our data suggest that IFGorb and IFGtri play different roles in processing ambiguous words, in line with the established distinction between retrieval and selection functions (Badre & Wagner, 2007). Our data are also compatible with the more general assertion that anterior IFG is specifically engaged by semantic processing while posterior IFG plays a more general role in various aspects of controlled language processing (Gough et al., 2005; Krieger-Redwood et al., 2015); and with functional and structural connectivity data indicating that IFGorb shows strong connectivity with anterior temporal regions linked specifically with semantic processing while more posterior regions (BA44/45) are also connected with frontoparietal networks involved in domain-general cognitive control (Jackson et al., 2016; Jung et al., 2016; Xiang et al., 2009).

### Anterior temporal lobe

The hub-and-spoke model holds that the ATL acts as a hub for integrating multi-modal information into conceptual representations, with the ventral portion of the ATL forming the heteromodal centre of this representational region (Lambon Ralph et al., 2017; Rice et al., 2015). The role of this region in comprehending homonyms has rarely been investigated; this is a significant lacuna because there are differing theoretical perspectives on the degree to which these conceptual representations should be sensitive to context (Binney et al., 2012; Hoffman et al., 2018; Lambon Ralph et al., 2017). We found that vATL exhibited distinct neural patterns for the same word depending on which of its meanings was currently relevant. This above-chance decoding is unlikely to reflect differences in processing effort between different meanings because this region is not sensitive to other experimental manipulations that varied in difficulty (i.e., it showed no univariate effects of relatedness or ambiguity). Instead, the results suggest that semantic representations coded in vATL are context-sensitive. This result supports the view that the hub uses recent linguistic context to shape its representations (Hoffman et al., 2018). This is advantageous because it allows the development of semantic representations to be informed by the statistical structure of natural language, which is a potentially valuable source of information about word meaning (Andrews et al., 2009; Landauer & Dumais, 1997; Mikolov et al., 2013).

How does this result fit with other theories that emphasise the context-independence of the hub? Such theories have typically defined context in terms of a top-down representation of the current task or goal, arguing that the ATL must be insensitive to these demands in order to acquire unbiased knowledge about the statistical structure of the environment (Binney et al., 2012; Lambon Ralph et al., 2017). Recent computational simulations support this view. By manipulating connectivity patterns in a connectionist hub-and-spoke model, Jackson et al. (2019) showed that semantic information was learned optimally when task representations were not allowed to influence activation in the model’s central conceptual hub. However, this goal-based context is a somewhat different idea to the bottom-up priming of meaning that we focus on in the present study. Indeed, our data suggest that ventral ATL *is* relatively insensitive to task requirements and current goals. Patterns in this region did not code for the semantic relatedness of trials, even when this was critical to the task being performed. Nor did it show any univariate effects of ambiguity or relatedness, indicating that the demand characteristics of different trial types did not influence its engagement. But despite this, its activation patterns did vary according to which homonym meaning was primed. Thus, the present data suggest that processing in ventral ATL is relatively “insulated” from top-down goal-related context, but at the same time is influenced by the bottom-up context provided by recent experience.

In contrast to ventral ATL, we found few effects in the more lateral portion of the ATL. This region showed no coding in MVPA analyses, which is surprising given its established role in verbal semantic cognition and in combinatorial semantic processing in particular (Baron & Osherson, 2011; Bemis & Pylkkänen, 2012; Humphries et al., 2006; Westerlund & Pylkkänen, 2014). Of course, there are a number of possible explanations for these null results. It may be that our region of interest did not cover the lateral regions most engaged by the task, or that it encompassed two different functional regions with distinct patterns of coding. At a univariate level, lateral ATL showed greater engagement for unambiguous words. This result is consistent with previous findings of greater engagement in this region for concepts that can be combined more easily (Hoffman et al., 2015; Teige et al., 2019).

### Angular gyrus

Angular gyrus presented a different pattern of effects to those in ATL and IFG. It displayed increased engagement for related trials (the opposite effect to IFG regions) and, unlike IFG regions, these effects were present in both the phonological and semantic tasks. Its patterns also coded for the presence or absence of a semantic relationship but, unlike IFGtri, this was true for the phonological as well as the semantic task. Before setting out our preferred interpretation of these findings, it is worth ruling out an alternative account. As a core element of the default mode network, AG frequently displays task-related deactivation in many cognitive domains, which increases with task difficulty (Humphreys et al., 2015; Mckiernan et al., 2003). This non-specific disengagement has been proposed to account for effects of semantic manipulations in some previous studies, rather than genuine involvement in semantic processing (Hoffman et al., 2015; Humphreys et al., 2015; Lambon Ralph et al., 2017). This was not the case here, however. In the semantic task, it was true that related trials were easier than unrelated trials (in terms of reaction time), which might have contributed to an effect of relatedness in AG. However, a relatedness effect was also observed in the phonological task, where there was no differences in behavioural performance.

As most effects in AG were observed independently of task, it seems that this region is not involved in goal-directed or controlled semantic processing. However, it does appear to be sensitive to the conceptual content of the stimuli, and in particular in the degree to which a coherent combinations of concepts is present. These results support the idea that AG is automatically engaged by the processing of coherent conceptual combinations (Davey et al., 2015; Humphreys & Lambon Ralph, 2014).

The precise role of AG in semantic processing is disputed. Some theories posit that AG acts as a semantic hub coding event-related or thematic semantic knowledge (Binder & Desai, 2011; Mirman et al., 2017; Schwartz et al., 2011). Others have suggested that AG serves as a short-term buffer for recent multimodal experience (Humphreys & Lambon Ralph, 2014). On the latter view, its function is not specific to semantic cognition but is required in some semantic tasks, particularly when context must be used to constrain semantic processing. Both of these accounts are consistent with our data. Related word pairs are more likely to activate representations of a coherent event than unrelated pairs, which could explain greater AG engagement on these trials. However, similar results might be expected if this region was involved in short-term maintenance of the target word in its particular context.

### Posterior middle temporal gyrus

Our final aim was to clarify the role of pMTG in processing semantic ambiguity. pMTG frequently shows increased engagement when homonyms are processed (Rodd et al., 2005; Rodd et al., 2015; Zempleni et al., 2007). In the present study, the effect of homonymy was not significant in our pMTG ROI (uncorrected *p* = 0.058), though whole-brain analysis did reveal greater activation for homonyms in this general anatomical region. Some studies have implicated pMTG in semantic control functions (Jefferies, 2013; Noonan et al., 2013; Whitney et al., 2011) while others have associated it with lexical-semantic representation (Bedny et al., 2008; Lau et al., 2008; Tyler et al., 2013). Our results are not wholly consistent with either view. Neural patterns in pMTG coded the semantic status of the trial, but only when this was task-relevant. In this sense, its response was similar to that of IFGtri, and suggests a general role in using semantic information to determine a behavioural response. However, unlike the IFG regions, engagement of pMTG was not increased for the more demanding semantically unrelated trials. In addition, pMTG activation patterns discriminated between the two meanings of homonyms. This might indicate either a role in representation of semantic knowledge or in controlling retrieval of such knowledge (akin to IFGorb). Overall, this complex set of findings is perhaps best explained in terms of a hybrid role for pMTG, whereby it is somewhat influenced by task demands but also by the nature of the semantic content being accessed. This fits well with other accounts claiming that pMTG is a functional nexus linking executive/semantic control networks with semantic representational regions including ATL and AG (Davey et al., 2015).

### Conclusions and future directions

The present study has revealed a complex set of responses to homonyms within the left-hemisphere semantic network. Ventral ATL showed systematic variation in activation patterns as a function of homonym meaning, suggesting that the semantic information coded in this region is sensitive to context. Activation in this area was similar across word and trial types, in line with representational system that processes stimulus information independently of task demands. In contrast, IFG regions showed variation in engagement as a function of task, indicating a goal-directed role in manipulation and evaluation of semantic information. However, we found that different subregions of IFG coded different forms of information about the stimuli, consistent with the idea that the more anterior IFGorb is engaged in top-down control over the retrieval of specific semantic information while IFGtri is resolving semantic competition in order to determine a behavioural response.

Importantly, we found that considering univariate and multivariate effects in combination provided important additional information about the functions of specific regions. For example, the MVPA analyses revealed very similar effects in ventral ATL and IFGorb: both regions decoded the currently-relevant meaning but not the semantic status of the trial, implicating them in coding and activating the specific semantic properties of the stimuli being presented. However, these regions dissociated in the univariate analyses, with only IFGorb showing greater engagement on unrelated trials (albeit only at an uncorrected statistical threshold). Changes in engagement are thought to reflect the degree to which a stimulus draws on a cognitive process supported by the region (Taylor et al., 2013). In this case, it appears that the process supported by IFGorb is more demanding when no semantic relationship is present, and we have argued that this process is likely to be controlled semantic retrieval. In contrast, the similar levels of activation for related/unrelated and for homonyms/unambiguous trials in ventral ATL suggests that processes supported by this region are engaged equally under all conditions. This is consistent with a more passive representational role which is engaged equally by comprehension under all circumstances.

What sort of representation is encoded by the ventral ATL? Connectionist models of semantic processing often posit that sentence processing involves the incremental formation of a gestalt representation that combines information about the word being currently processed with its prior context (Elman, 1990; Rabovsky et al., 2018; St. John, 1992). On this view, the pattern of semantic activation elicited by a particular word is not constant, but varies depending on the exact context in which appears (Cruse, 1986; Hoffman & Woollams, 2015; Landauer, 2001). Homonyms are subject to particularly large shifts in representation because of context has such a large effect on their interpretation. For our study, this approach predicts some variability in representation even within trials that prime the same meaning (e.g., *BARK* in the context of *tree* is similar but not identical to *BARK* in the context of *wood*), but a greater distinction between these trials and those in which the opposing meaning is primed.

An alternative view is that the meanings of a homonym are represented as two different entries in a lexicon, with the more appropriate entry activated on each occasion (Duffy et al., 1988; Kellas et al., 1988). This view might envision no difference between the representations of *BARK* in the context of *tree* vs. *wood*, though both would differ from *BARK* in the context of *dog*. Our data cannot adjudicate between these different possibilities. Indeed, because there was only a two-second interval between prime and target, we cannot rule out the possibility that neural activity elicited at the point of prime activation also contributed to the decoding we observed. To distinguish between these various options, future studies could vary the delay between prime and target to better separate the neural correlates of each. This could involve separating prime and target with multiple intervening trials, since priming of homonym meanings has been shown to persist over long intervals (Rodd et al., 2013).

While multiple regions showed meaning-specific coding for homonyms in the semantic task, no regions showed this effect when participants made phonological decisions about the words. Does this result indicate that the brain does not disambiguate homonyms under these conditions? Not necessarily. All of our ROIs showed lower levels of engagement for the phonological task relative to the semantic task. Thus, it seems that when participants are focused on performing phonological judgements, in which semantic information is unhelpful, neural resources are directed away from the semantic system. One corollary of this is that signal in these regions is much weaker, and may have been insufficient for the classifier to predict meaning at above-chance level. In other words, it is possible that there were still subtle activation shifts as a function of meaning during phonological processing, but not at a level that we were able to reliably detect with fMRI. Alternatively, it may be that minimal semantic processing occurs under these conditions. One final possibility is that disambiguating neural information is briefly activated upon stimulus processing but is not sustained. The temporal resolution of fMRI is such that brief changes in activity are less likely to be detected. Studies using multivariate decoding methods applied to EEG or MEG data may be valuable in providing more information about the time-course of semantic activation during homonym comprehension.

Finally, we acknowledge that our experimental paradigm, using pairs of individual words, is far removed from comprehension in more naturalistic language contexts. Each of our homonyms was also repeated eight times during each task, requiring repeated switching between interpretations that is not representative of natural language processing. These design choices were necessary in order to obtain a sufficient number of trials for MVPA analysis. We are reassured by the fact that our univariate analysis revealed similar effects of ambiguity as other fMRI studies that have presented ambiguous words presented in natural sentences (Mason & Just, 2007; Rodd et al., 2005; Zempleni et al., 2007). Nevertheless, it remains to be seen whether distinct activation patterns elicited by different meanings would hold when people process more naturalistic sentence-level stimuli. Indeed, the vast majority of studies that have so far used MVPA to investigate semantic representation have focused on the single-word level. The extension of these methods to the sentence level and beyond will be critical in gaining a more complete understanding of the neural basis of semantic representation.

## Acknowledgements

The study was funded by grants from the Carnegie Trust for the Universities of Scotland (70743) and Moray Endowment Fund. PH was also supported by the University of Edinburgh Centre for Cognitive Ageing and Cognitive Epidemiology, part of the cross council Lifelong Health and Wellbeing Initiative (MR/K026992/1). Funding from the Biotechnology and Biological Sciences Research Council (BBSRC) and Medical Research Council (MRC) is gratefully acknowledged.

## Supplementary Materials

**Supplementary Table 1:**
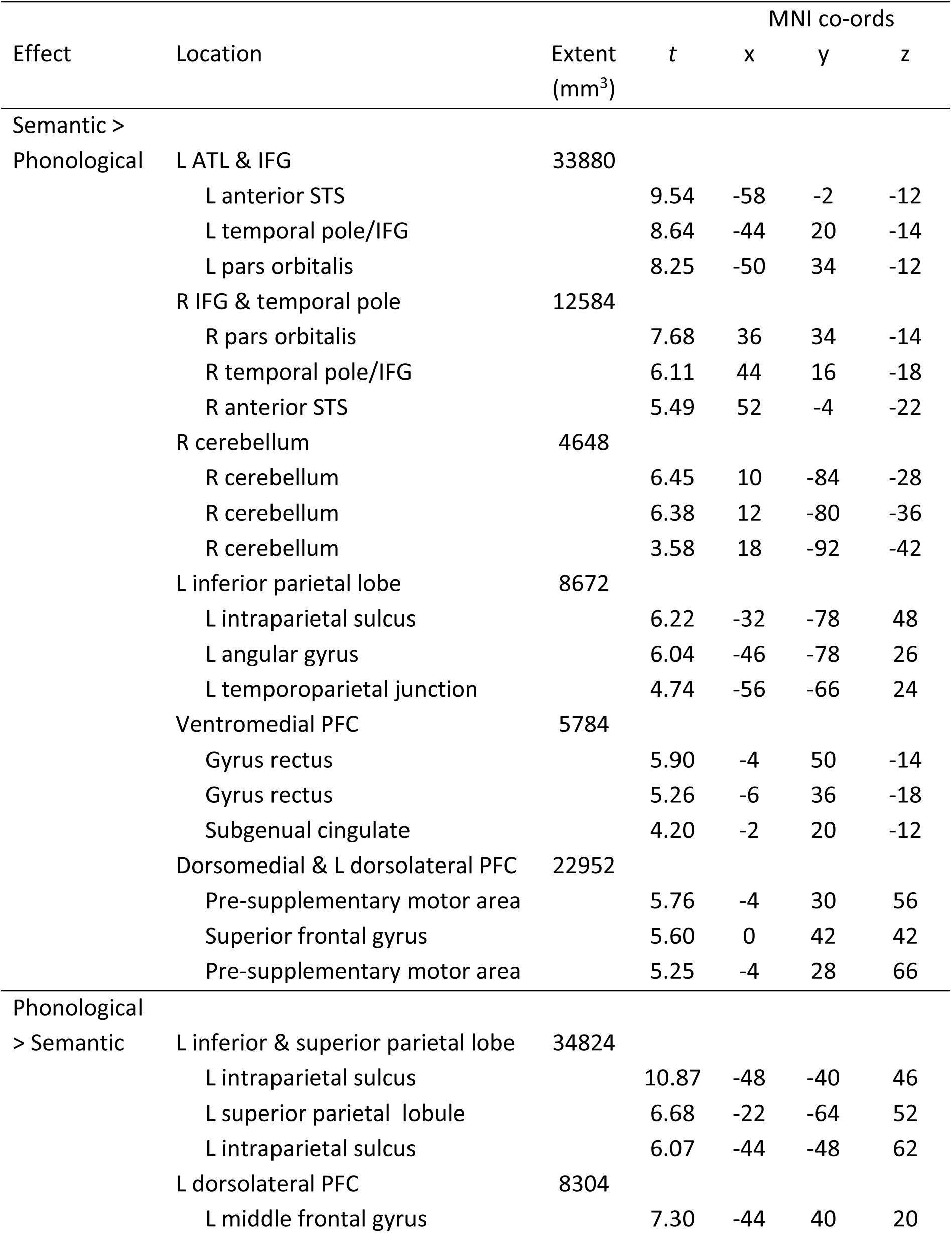
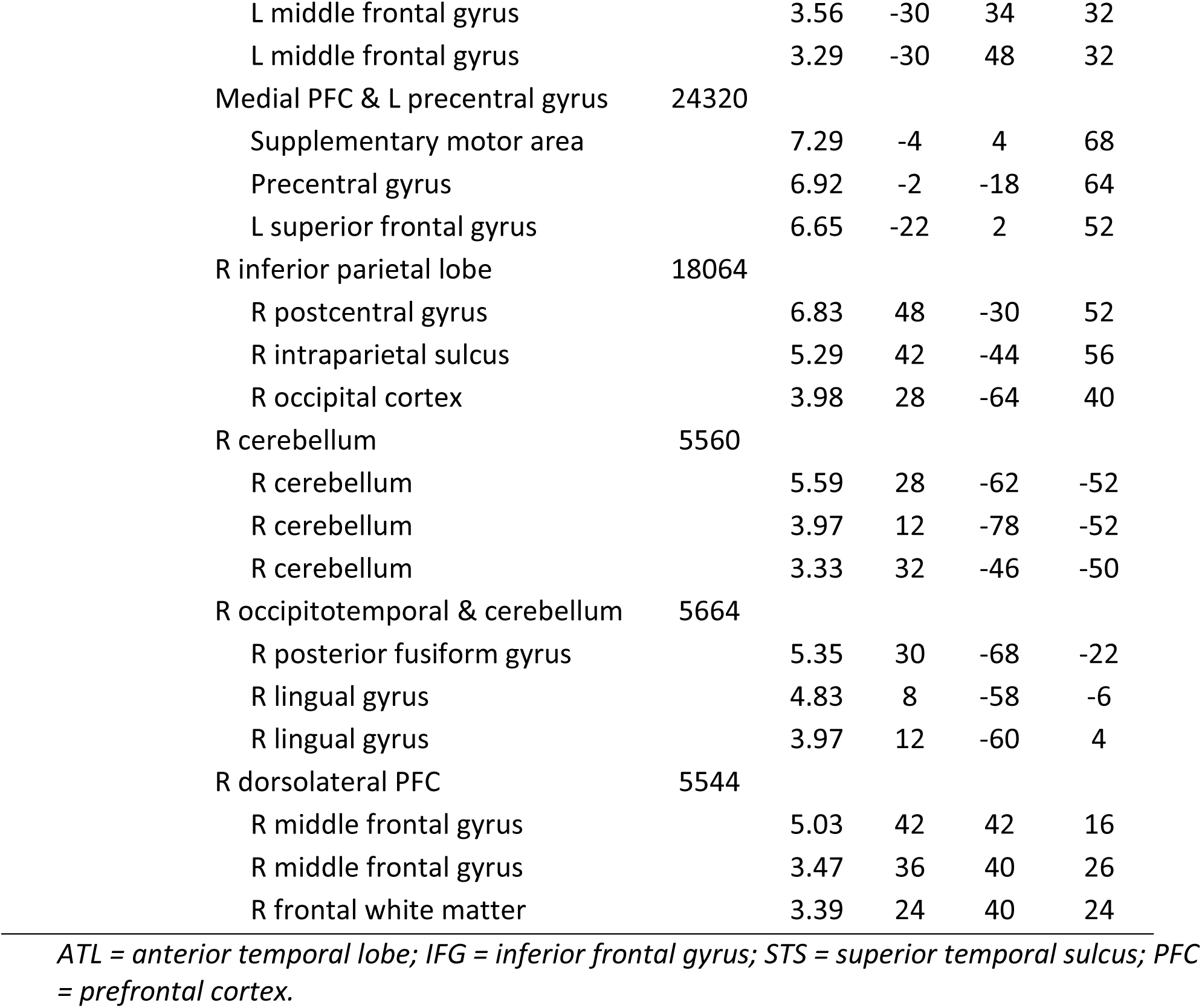
Peak activation co-ordinates for univariate contrasts of task

| Effect | Location | Extent<br>(mm<sup>3</sup>) | t | MNI co-ords |  |  |
| --- | --- | --- | --- | --- | --- | --- |
|  |  |  |  | x | y | z |
| Semantic > |  |  |  |  |  |  |
| Phonological | L ATL & IFG | 33880 |  |  |  |  |
|  | L anterior STS |  | 9.54 | -58 | -2 | -12 |
|  | L temporal pole/IFG |  | 8.64 | -44 | 20 | -14 |
|  | L pars orbitalis |  | 8.25 | -50 | 34 | -12 |
|  | R IFG & temporal pole | 12584 |  |  |  |  |
|  | R pars orbitalis |  | 7.68 | 36 | 34 | -14 |
|  | R temporal pole/IFG |  | 6.11 | 44 | 16 | -18 |
|  | R anterior STS |  | 5.49 | 52 | -4 | -22 |
|  | R cerebellum | 4648 |  |  |  |  |
|  | R cerebellum |  | 6.45 | 10 | -84 | -28 |
|  | R cerebellum |  | 6.38 | 12 | -80 | -36 |
|  | R cerebellum |  | 3.58 | 18 | -92 | -42 |
|  | L inferior parietal lobe | 8672 |  |  |  |  |
|  | L intraparietal sulcus |  | 6.22 | -32 | -78 | 48 |
|  | L angular gyrus |  | 6.04 | -46 | -78 | 26 |
|  | L temporoparietal junction |  | 4.74 | -56 | -66 | 24 |
|  | Ventromedial PFC | 5784 |  |  |  |  |
|  | Gyrus rectus |  | 5.90 | -4 | 50 | -14 |
|  | Gyrus rectus |  | 5.26 | -6 | 36 | -18 |
|  | Subgenual cingulate |  | 4.20 | -2 | 20 | -12 |
|  | Dorsomedial & L dorsolateral PFC | 22952 |  |  |  |  |
|  | Pre-supplementary motor area |  | 5.76 | -4 | 30 | 56 |
|  | Superior frontal gyrus |  | 5.60 | 0 | 42 | 42 |
|  | Pre-supplementary motor area |  | 5.25 | -4 | 28 | 66 |
| Phonological |  |  |  |  |  |  |
| > Semantic | L inferior & superior parietal lobe | 34824 |  |  |  |  |
|  | L intraparietal sulcus |  | 10.87 | -48 | -40 | 46 |
|  | L superior parietal lobule |  | 6.68 | -22 | -64 | 52 |
|  | L intraparietal sulcus |  | 6.07 | -44 | -48 | 62 |
|  | L dorsolateral PFC | 8304 |  |  |  |  |
|  | L middle frontal gyrus |  | 7.30 | -44 | 40 | 20 |

|  |  |  |  |  |  |
| --- | --- | --- | --- | --- | --- |
| L middle frontal gyrus |  | 3.56 | -30 | 34 | 32 |
| L middle frontal gyrus |  | 3.29 | -30 | 48 | 32 |
| Medial PFC & L precentral gyrus | 24320 |  |  |  |  |
| Supplementary motor area |  | 7.29 | -4 | 4 | 68 |
| Precentral gyrus |  | 6.92 | -2 | -18 | 64 |
| L superior frontal gyrus |  | 6.65 | -22 | 2 | 52 |
| R inferior parietal lobe | 18064 |  |  |  |  |
| R postcentral gyrus |  | 6.83 | 48 | -30 | 52 |
| R intraparietal sulcus |  | 5.29 | 42 | -44 | 56 |
| R occipital cortex |  | 3.98 | 28 | -64 | 40 |
| R cerebellum | 5560 |  |  |  |  |
| R cerebellum |  | 5.59 | 28 | -62 | -52 |
| R cerebellum |  | 3.97 | 12 | -78 | -52 |
| R cerebellum |  | 3.33 | 32 | -46 | -50 |
| R occipitotemporal & cerebellum | 5664 |  |  |  |  |
| R posterior fusiform gyrus |  | 5.35 | 30 | -68 | -22 |
| R lingual gyrus |  | 4.83 | 8 | -58 | -6 |
| R lingual gyrus |  | 3.97 | 12 | -60 | 4 |
| R dorsolateral PFC | 5544 |  |  |  |  |
| R middle frontal gyrus |  | 5.03 | 42 | 42 | 16 |
| R middle frontal gyrus |  | 3.47 | 36 | 40 | 26 |
| R frontal white matter |  | 3.39 | 24 | 40 | 24 |

*ATL = anterior temporal lobe; IFG = inferior frontal gyrus; STS = superior temporal sulcus; PFC = prefrontal cortex.*

**Supplementary Table 2:**
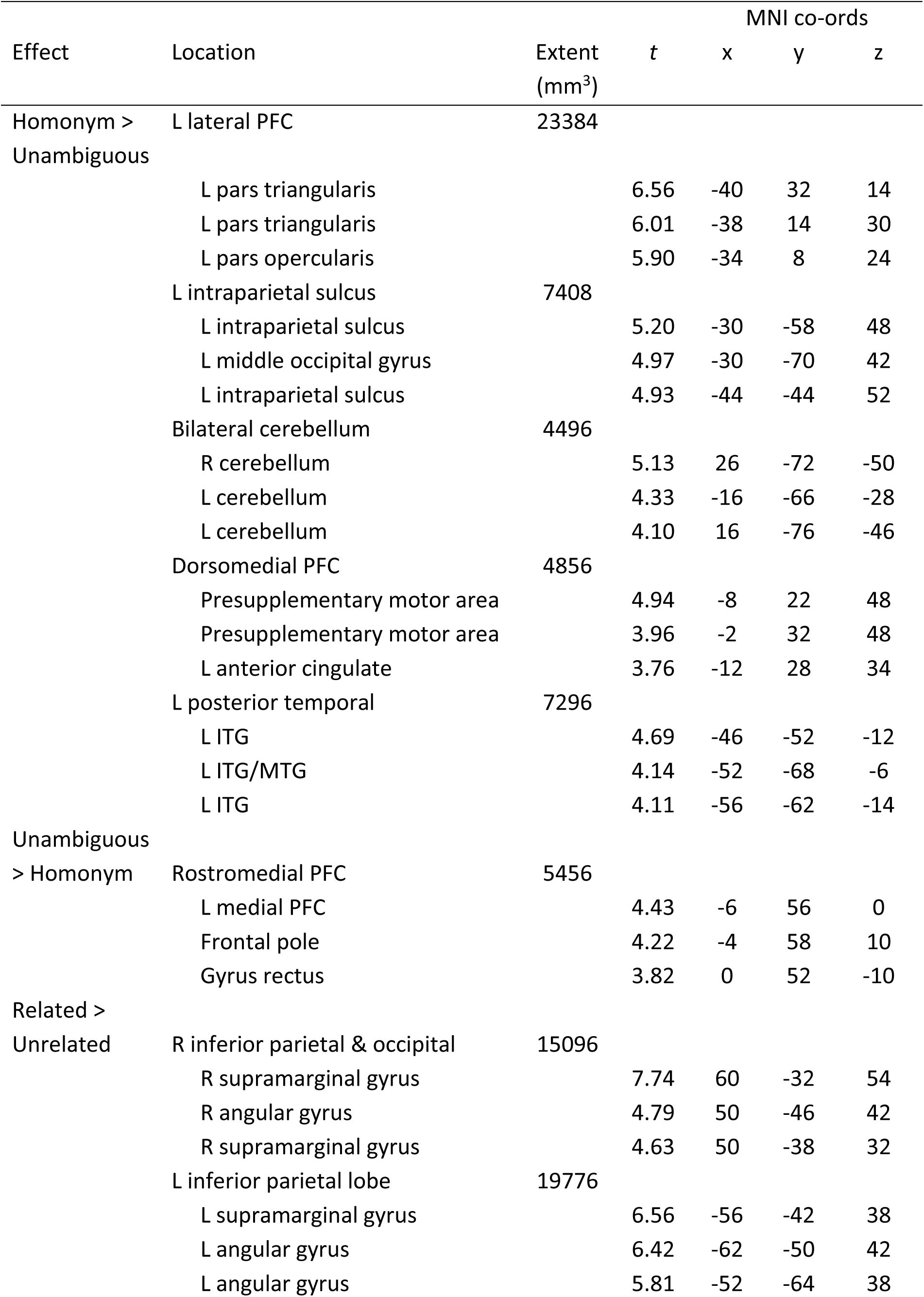
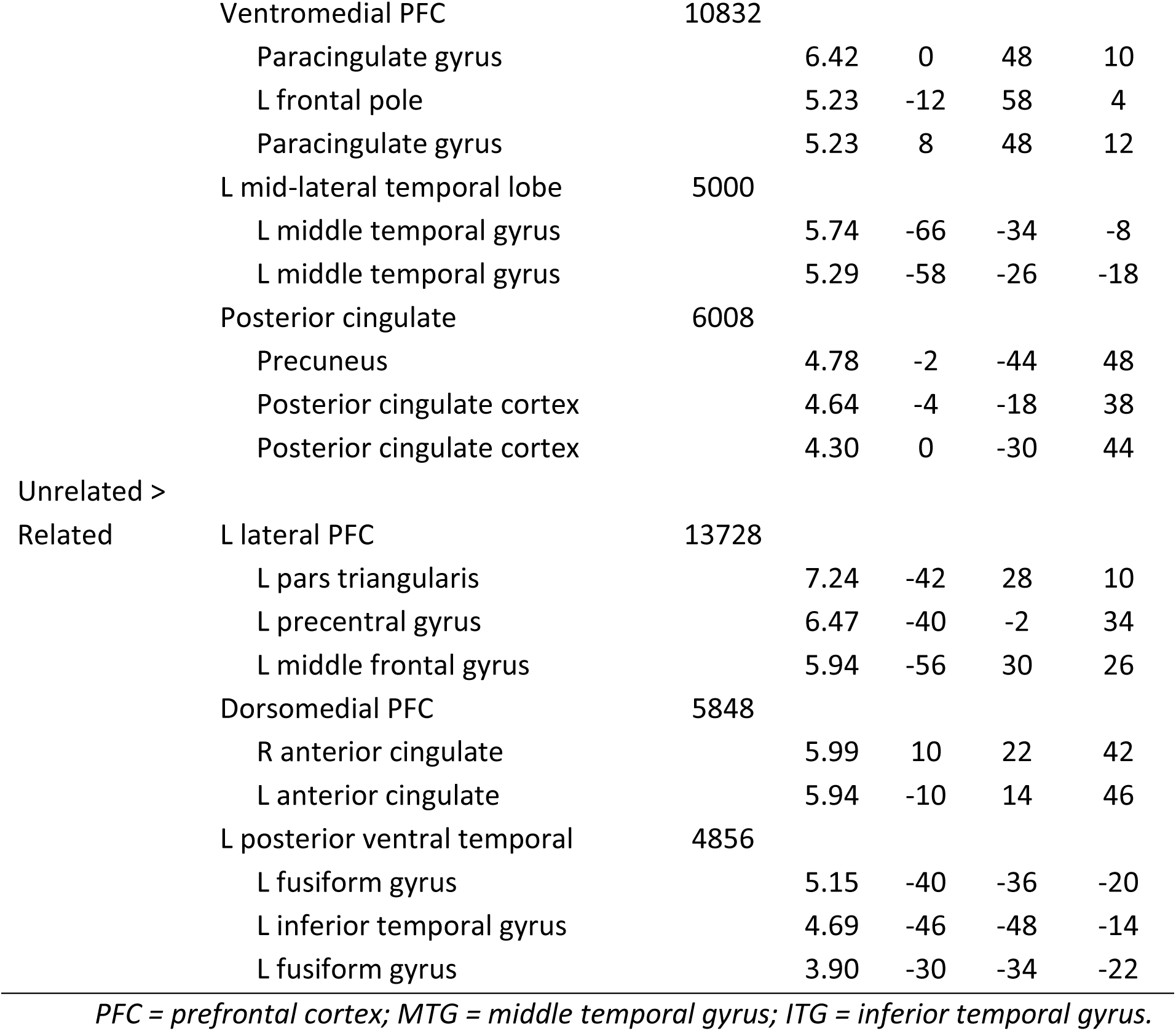
Peak activation co-ordinates for univariate contrasts in the semantic task

**Supplementary Figure 1:**
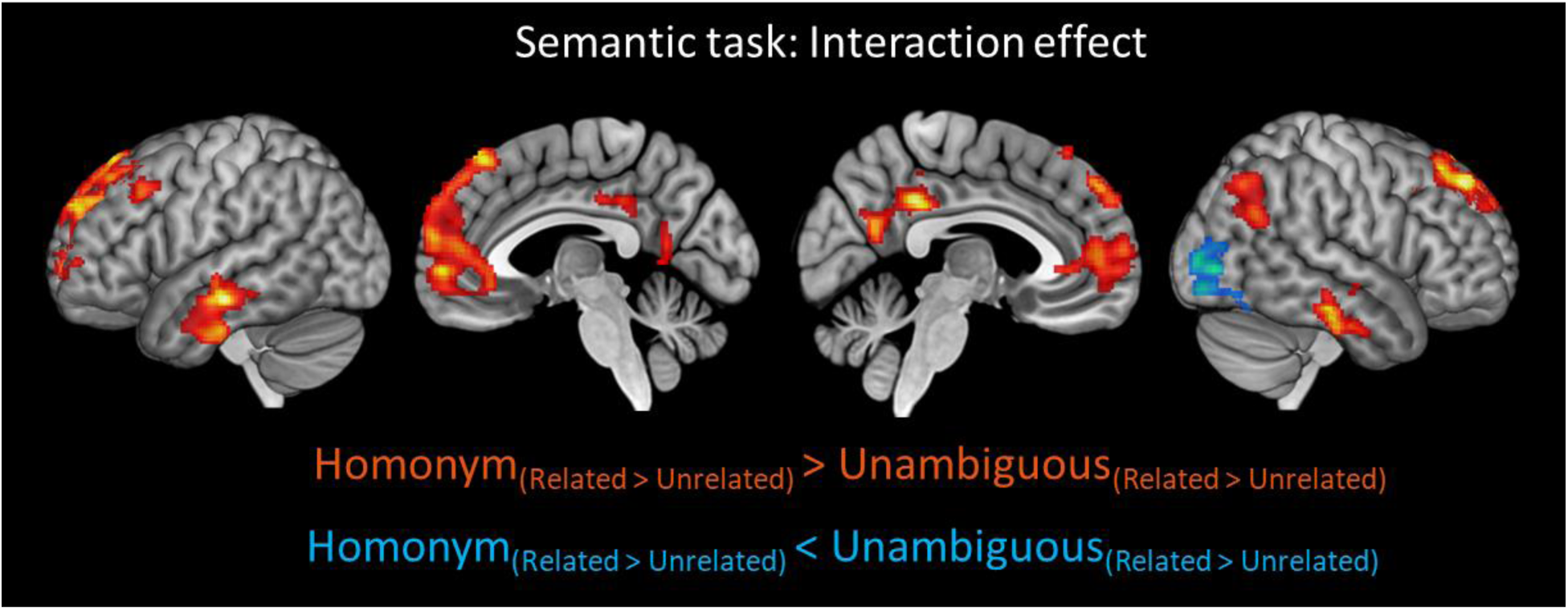
Whole-brain univariate activation contrasts for the interaction effect in the semantic task. Images are shown at a voxelwise threshold of p<0.005, corrected for multiple comparisons at the cluster level.

**Supplementary Figure 2:**
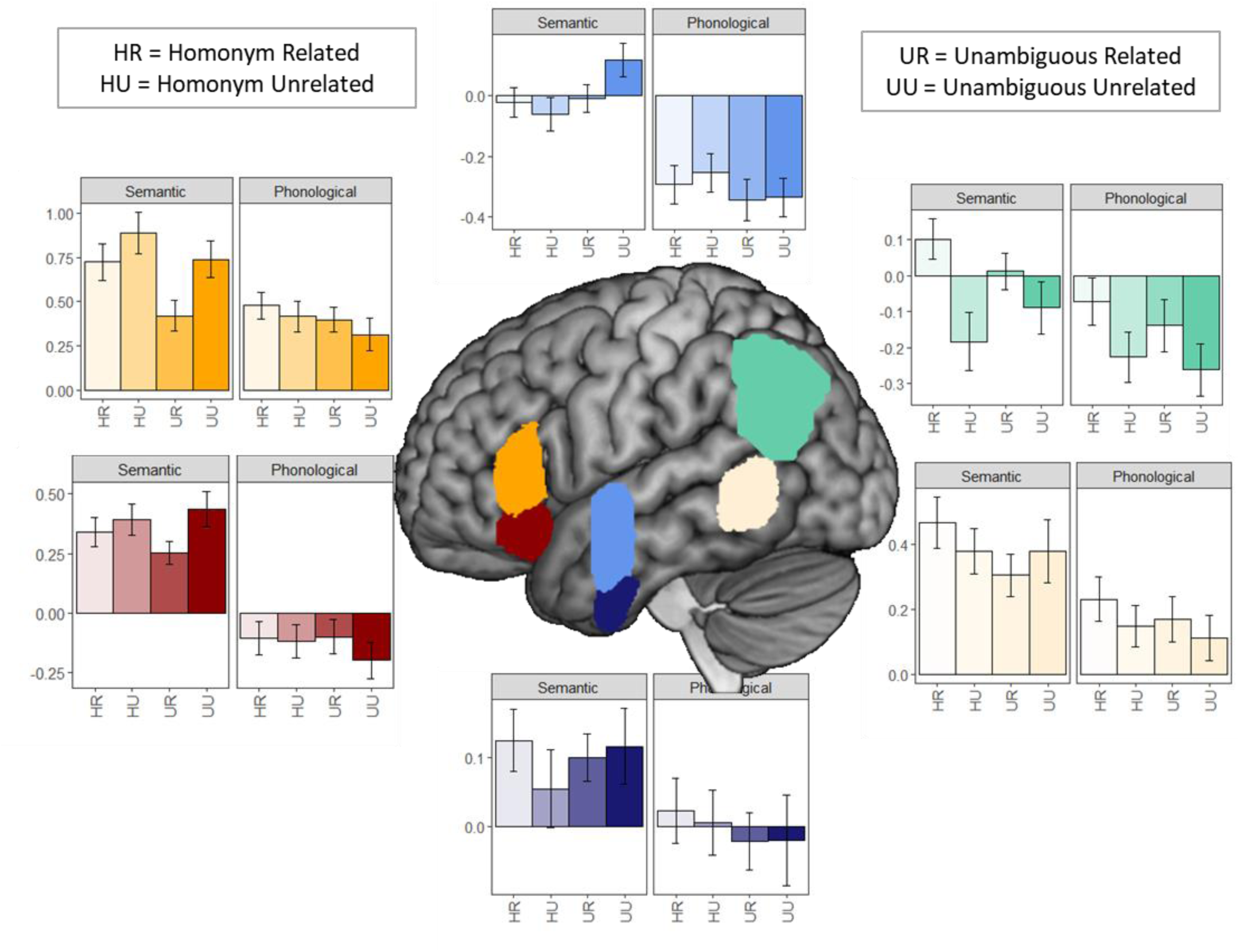
Activation in each region of interest as a function of condition. Effect size is shown relative to implicit resting baseline. Bars indicate between-subjects SEM.

**Supplementary Figure 3:**
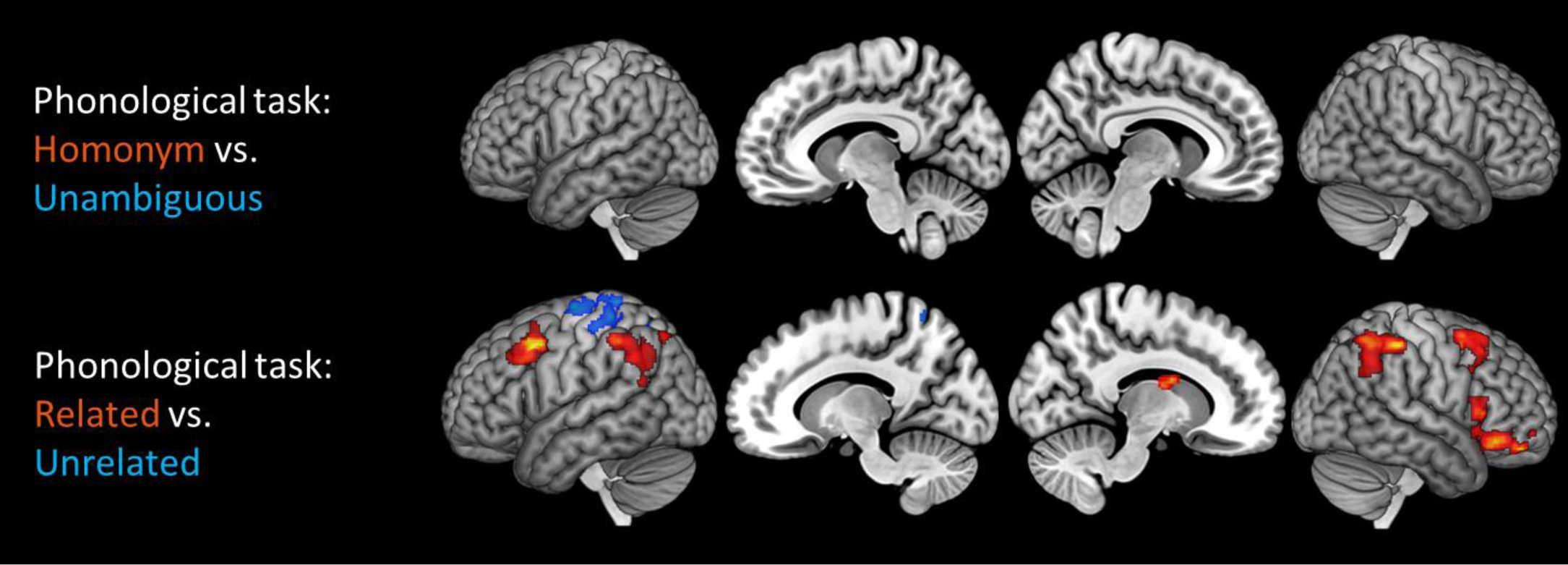
Whole-brain univariate activation contrasts for the phonological task. Images are shown at a voxelwise threshold of p<0.005, corrected for multiple comparisons at the cluster level.

